# Homozygous kinase-dead *Csf1r* mutation in outbred mice reveals essential and redundant functions of tissue resident macrophages

**DOI:** 10.1101/2025.07.08.663807

**Authors:** Sebastien Jacquelin, Emma Maxwell, Isis Taylor, Emma Green, Conan O’Brien, Ginell Ranpura, Jintao Guo, Fathima Nooru-Mohamed, Yajun Liu, Sahar Keshvari, Stephen Huang, Emily Cooper, Steven W. Lane, Cameron Flegg, Kamil Sokolowski, Allison R. Pettit, David A. Hume, Katharine M. Irvine

**Author notes:** Joint senior and corresponding authors.

## Abstract

The proliferation, differentiation and survival of cells of the macrophage lineage depends on signals from the macrophage colony-stimulating factor receptor (CSF1R). On a C57BL/6J background homozygous kinase-dead *Csf1r* mutation (*Csf1r*^E631K/E631K^ - E631K^m/m^) causes perinatal lethality. Here we demonstrate that E631K^m/m^ mice on a mixed genetic background (C57 x BALB/c F2) are osteopetrotic and growth retarded but viable as adults with no other gross developmental deficits. They lack osteoclasts, microglia and most peripheral tissue resident macrophages and exhibit perturbed hematopoiesis. Although CD169^+^ tissue resident macrophages in bone marrow are considered an essential component of the hematopoietic niche, CD169 is undetectable in E631K^m/m^ marrow and F4/80^+^ macrophages are depleted. These changes are associated with expansion of mature and immature granulocytes and reduced B cells, whereas monocytes and stem and progenitor populations are unaffected as a proportion of total cells. Erythropoiesis in bone marrow is maintained in E631K^m/m^ mice, associated with a residual population of CSF1R-independent CD169^-ve^/F4/80^+^ macrophages. Nevertheless, splenic extramedullary hematopoiesis in E631K^m/m^ mice indicates a degree of bone marrow insufficiency. Red pulp macrophages are retained but CD169^+^ marginal metallophil macrophages are absent and CD209b^+^ (SIGNR1) macrophages are present but disorganized. Circulating white blood cell count is unchanged in E631K^m/m^ mice, but the proportion of neutrophils is greatly increased whilst B cells and monocytes are reduced. This novel model reveals the essential roles of CSF1R-dependent macrophages in hematopoiesis and demonstrates that many developmental and homeostatic functions attributed to murine resident tissue macrophages are redundant and/or specific to inbred mouse strains.

## Introduction

Colony stimulating factor 1 receptor (CSF1R) signalling in response to its two alternative ligands, CSF1 and IL34, controls survival, proliferation, and differentiation of mononuclear phagocyte populations throughout the body ^1,2^. Upon ligand binding, CSF1R dimerization and autophosphorylation generates phosphotyrosine docking sites for multiple downstream effector pathways ^1,3^. Biallelic recessive loss-of-function mutations in rat ^4–7^ and human ^8,9^ *CSF1R* lead to the loss of most tissue mononuclear phagocyte populations including osteoclasts and microglia. The associated postnatal developmental abnormalities include osteopetrosis, hydrocephalus and leukoencephalopathy and early mortality. Homozygous *Csf1r* knockout (*Csf1rko*) in mice on a mixed genetic background also led to macrophage deficiency and osteopetrosis that resembled the phenotype seen in a natural mutation of the *Csf1* gene ^10^. The same mutation on defined inbred backgrounds (C57BL/6J, FVB/J) leads to perinatal or early postnatal lethality ^11–14^. Consequently, *Csf1rko* mice have not been widely studied.

Heterozygous mutations that ablate CSF1R kinase activity are associated with adult-onset leukoencephalopathy ^15–19^. When expressed in a factor-dependent cell line, mutant CSF1R trafficked to the cell surface and bound and internalised CSF1 but failed to support cell survival or proliferation ^20^, consistent with evidence that these mutations abrogate CSF1R kinase activity ^21^. We generated a transgenic mouse line with a disease-associated *Csf1r* mutation (E631K, E633K in humans) ^21^. Although they lacked macrophages, *Csf1r*^E631K/E631K^ embryos at 12.5 dpc appeared indistinguishable from littermates ^22^ but the few viable homozygous pups born were severely growth-retarded and succumbed to hydrocephalus ^23^. To mitigate the perinatal lethality, we generated *Csf1r*^E631K/E631K^ mice on a C57BL/6J x BALB/c F2 background. These mice provide a unique model to dissect the functions of CSF1R signalling in monocyte-macrophage differentiation and CSF1R-dependent macrophages in hematopoiesis, homeostasis and postnatal development.

## Results

### Csf1r^E631K/E631K^ outbred mice are viable

The *Csf1r*^E631K^ mutant allele on C57BL/6J background was crossed to the *Csf1r-*EGFP reporter transgenic line ^24^. Heterozygous matings of *Csf1r*^E631K/+^ mice on this background produced no viable homozygous offspring ^23^. We crossed *Csf1r*^E631K/+^/*Csf1r-*EGFP^+^ male mice with BALB/c female mice and intercrossed the F1 progeny (**Figure 1A**). The genotype frequencies of F2 WT, *Csf1r*^E631K/+^ and *Csf1r*^E631K/E631K^ (E631K^m/m^) at weaning (**Figure 1B**) indicate that perinatal lethality in E631K^m/m^ mice is not a simple Mendelian trait but rather is overcome entirely on the mixed genetic background. E631K^m/m^ pups were not morphologically distinguishable from littermates at birth but postnatal somatic growth was impaired. Growth plateaued at 6-8 weeks of age; mutant mice did not exhibit the continued growth associated with sexual maturity in WT (**Figure 1C)**. A subset of males failed to thrive postweaning and required euthanasia whereas females were healthy up to the oldest analysed at 23 weeks.

**Figure 1.**
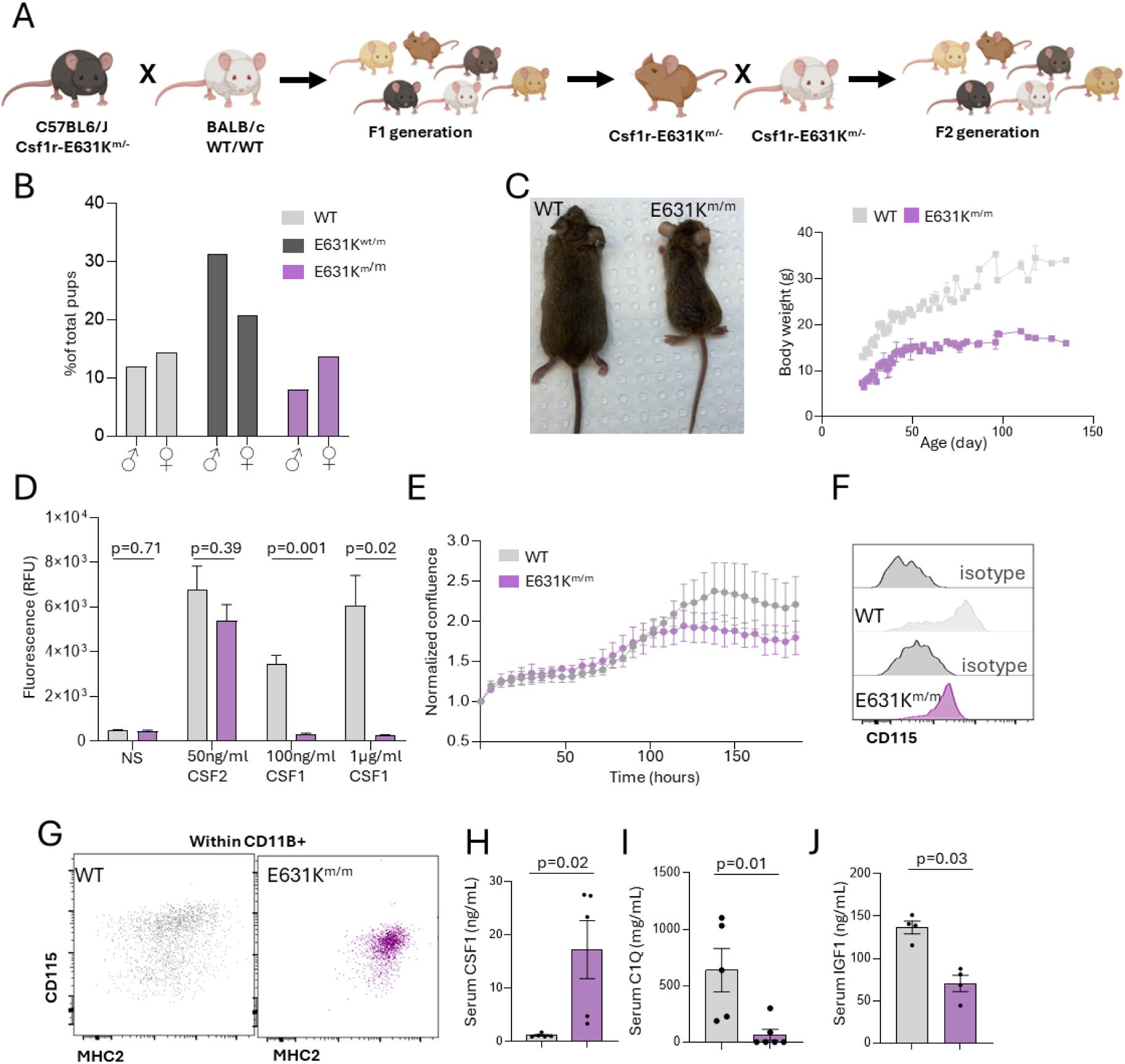
Generation and characterisation of outbred Csf1r E631K^m/m^ mutant Mice. (A) Summary of mouse breeding. First generation (F1) *Csf1r*^E631K/+^ mice from C57Bl6/J/ *Csf1r* ^E631K/+^ x BALB/c matings were intercrossed to produce F2 progeny including WT, heterozygous and homozygous mutant mice on a mixed genetic background (B) Proportions of WT, *Csf1r* ^E631K/+^ and E631K^m/m^ F2 pups at weaning, from a total of 133 pups. Differences in genotype frequency between male and female are not significant (C) Adult E631K^m/m^ mice are approximately half the size of WT littermates at the same age (10 weeks in image shown). Panel at right shows the combined post weaning growth curve (average body weight) for male and female WT and E631K^m/m^ mice. (D) Bone marrow cells from WT or E631K^m/m^ mice were cultured in CSF1 or CSF2 and metabolic activity after 6 days was measured using a resazurin assay (E) Bone marrow cells from WT or E631K^m/m^ mice were cultured in 50 ng/ml CSF2 for 7 days in an Incucyte with confluency measurement every 6 hours. Confluence was measured in quadruplicate for four mice per genotype. Data show means and standard deviation normalized to 0 hours. (F,G) BMM derived from WT and E631K^m/m^ cultured in CSF2 were analyzed by Flow Cytometry. Panels show typical profiles for CD115, CD11b and MHC2. CSF1 (J), C1Q (K) and IGF1 (L) were measured by ELISA in sera from adult (8-12 week) old WT and E631K^m/m^ female mice. Each point is an individual animal. Data are presented as mean and standard deviation. *p* values were determined by unpaired two-tailed t-test.

### The Csf1r^E631K/E631K^ mutation abrogates CSF1R signalling

We first sought to verify that E631K^m/m^ mice represent a functional CSF1R knockout. E631K^m/m^ bone marrow cells were unresponsive to CSF1 simulation *in vitro* whereas the proliferative response to CSF2 (GM-CSF) was not significantly reduced compared to WT (**Figure 1D,E)**. Surface CSF1R (CD115) was expressed on bone marrow-derived macrophages (BMM) derived by cultivation in CSF2, albeit at lower levels in E631K^m/m^ compared to WT mice (**Figure 1F**). Cultivation of bone marrow in CSF2 generates a mixed population of antigen-presenting cells differing in expression of class II MHC (MHC2) ^25^. CSF2-BMM from E631K^m/m^ mice were strongly polarised towards high MHC2 expression suggesting that endogenous CSF1/CSF1R signals modulate the response to CSF2 (**Figure 1G**).

We previously identified several serum markers of macrophage deficiency in *Csf1rko* rats ^4,26^. CSF1R^+^ macrophages regulate the levels of circulating CSF1 by receptor-mediated endocytosis ^27^. CSF1 was greatly elevated in E631K^m/m^ mouse serum (**Figure 1H).** Conversely the reduction in circulating complement C1Q and insulin-like growth factor 1 (IGF1) noted in *Csf1rko* rats was conserved in E631K^m/m^ mice (**Figure 1I,J).**

### The Csf1r^E631K/E631K^ mutation leads to osteopetrosis

Micro CT analysis of tibia revealed a striking increase in trabecular number and concomitant decrease in spacing in E631K^m/m^ mice (**Figure S1A,B**) associated with almost complete loss of tartrate resistant acid phosphatase (TRAP)^+^ osteoclasts (**Figure S1C**). Micro CT analysis of the jaw confirmed the presence of teeth and the failure of eruption of the incisors (**Supp-movie 1-2)**. The loss of osteoclasts and hematopoietic marrow in the vicinity of the metaphysis of long bones was also evident by combined staining for F4/80 and CD71 (transferrin receptor) (**Figure S1D**). CD71 staining also revealed the continued cell proliferation associated with the epiphysis in E631K^m/m^ mice (**Figure S1E**). WT bone marrow cells grown in CSF1 responded to RANK ligand to generate TRAP^+^ osteoclasts. By contrast, although CSF2 directed the production of BMM as shown in Figure 1, cells grown in CSF2 were unresponsive to RANKL regardless of genotype even if CSF1 was also present (**Figure S1F**).

**Figure S1.**
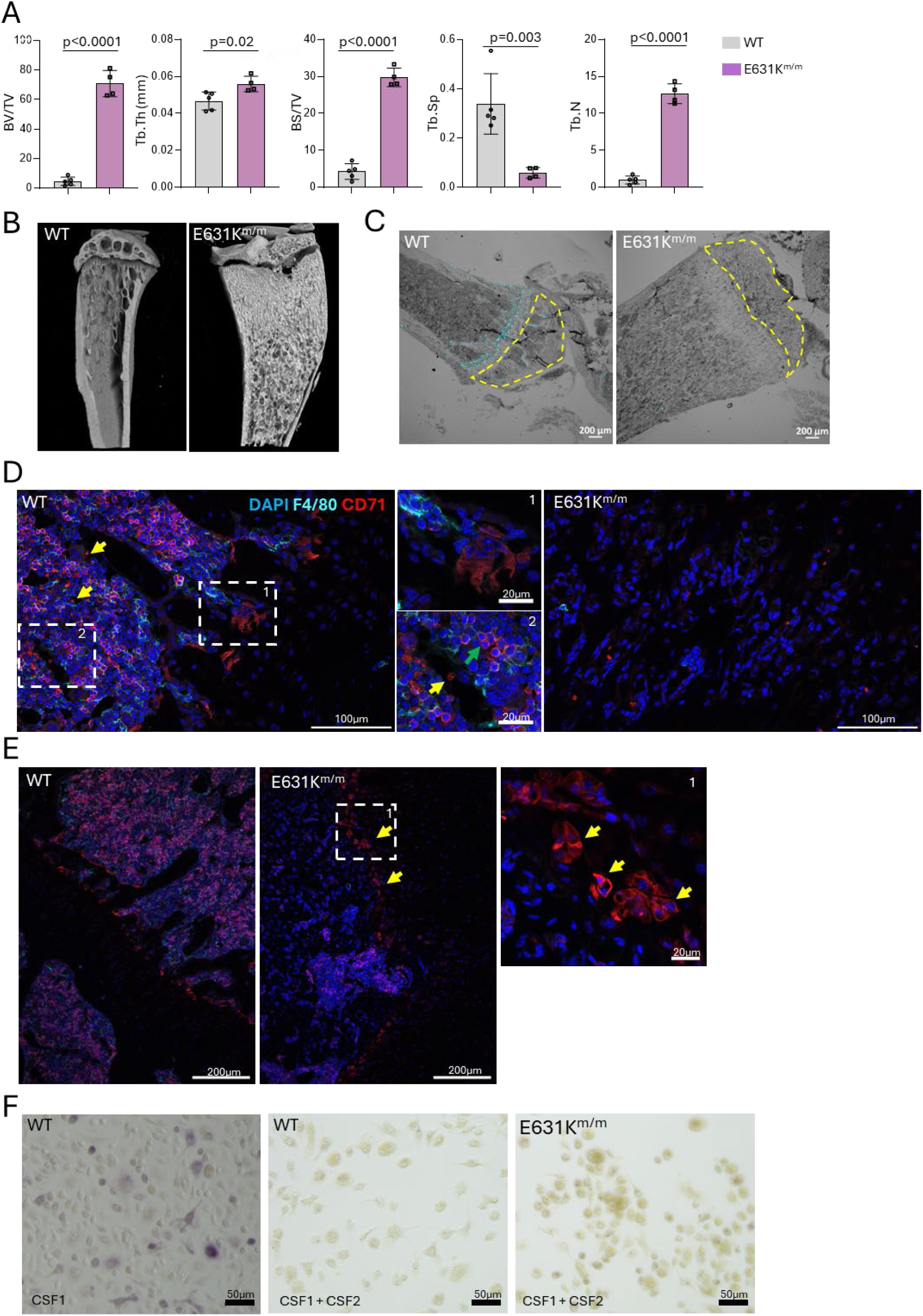
Osteopetrosis and osteoclast deficiency in E631K^m/m^ mice. (A) Quantitation of bone volume/trabecular volume (BV/TV), trabecular thickness (Tb.Th), bone surface/total volume (BS/TV), trabecular space (Tb.Sp) and trabecular number (Tb.N.) from microCT imaging of the tibial metaphysis in WT and E631K^m/m^ female mice at 8-12 weeks of age. Each point is an individual animal. Data are shown as mean and standard deviation. *p* values were determined by unpaired two-tailed t-test. (B) Representative 3D reconstruction of tibia from adult WT and E631K^m/m^ mice. (C) Sections of tibia from adult WT and E631K^m/m^ mice stained for tartrate resistant acid phosphatase (TRAP) (cyan) projected onto a bright field image of the same section. Note the presence of marrow within the diaphysis (yellow outline) in WT and expansion of the growth plate in the E631K^m/m^ mice. (D) Representative images of localisation of F4/80 and CD71 in diaphysis of adult WT and E631K^m/m^ mice. Insets in WT highlight CD71 in multinucleated osteoclasts, erythroblastic islands (green arrows) and reticulocytes (yellow arrows). (E) Representative images of localisation of F4/80 and CD71 in epiphysis of adult WT and E631K^m/m^ mice. Arrows and Insets highlight the presence of CD71^+^ cells on the surface of the growth plate in mutant mice. (E) Bone marrow cells from WT and E631K^m/m^ mice were cultured in CSF1 or CSF1+CSF2, treated with RANKL and stained for TRAP. Representative images are shown.

### E631K^m/m^ mice are globally macrophage deficient

Wholemount imaging of the *Csf1r-*EGFP reporter and/or immunohistochemical staining for F4/80 in adult E631K^m/m^ mice revealed almost complete loss of macrophages in most organs, including intercostal and heart muscle, subcutaneous and perigonadal white adipose tissue (WAT), endocrine and exocrine pancreas, adrenal and salivary glands, testis, bladder, kidney and ovary. Other tissues, including liver, lung, uterus and the gastrointestinal tract retained macrophage populations, albeit reduced (**Figure 2A-E**). In the uterus, imunolocalisation of F4/80 indicated the loss of stellate macrophage populations whereas monocyte-like cells were retained in E631K^m/m^ mice (**Figure 2B**). In the liver, whole mount imaging (**Figure 2D**) confirmed the absence of the CSF1R-dependent subcapsular macrophage population ^28^ in E631K^m/m^ mice. Residual F4/80^+^ macrophages in the parenchyma resembled Kupffer cells and were located within the sinusoids (**Figure 2D**). The abundant interstitial macrophages of the lung periphery were also evident in whole mounts from WT and absent in E631K^m/m^ mice (**Figure 2D**). In the intestine, the mucosa appeared thinner and macroscopic Peyer’s patches were less evident in E631K^m/m^ mice (**Figure S2A**). However, regular lymphoid aggregates were detected and villus architecture appeared normal (**Figure S2B**). The lamina propria and muscularis of small and large intestine contain an abundant population of stellate F4/80^+^ macrophages^29^. Muscularis macrophages can be imaged in whole mount using the *Csf1r*-EGFP transgene and were absent in E631K^m/m^ mice (see colon image in **Figure 2A**). However, F4/80^+^ cells were present in the submucosa and lamina propria (**Figure S2B**). At higher magnification, the cellularity of the lamina propria appeared reduced in the mutant mice and the residual F4/80^+^ cells had a more rounded monocyte-like morphology (**Figure S2C, D**). Histochemical staining indicated that the Paneth cells and goblet cells were not impacted (**Figure S2E**). Ki67 staining was similar between genotypes, indicating that intestinal epithelial cell turnover and replacement was not affected by macrophage deficiency (**Figure S2F**). Analysis of peritoneal lavage showed selective loss of F4/80^Hi^ resident peritoneal macrophages in E631K^m/m^ mice whereas the small peritoneal macrophages were retained and granulocytes were increased (**Figure S2G-I**).

**Figure 2.**
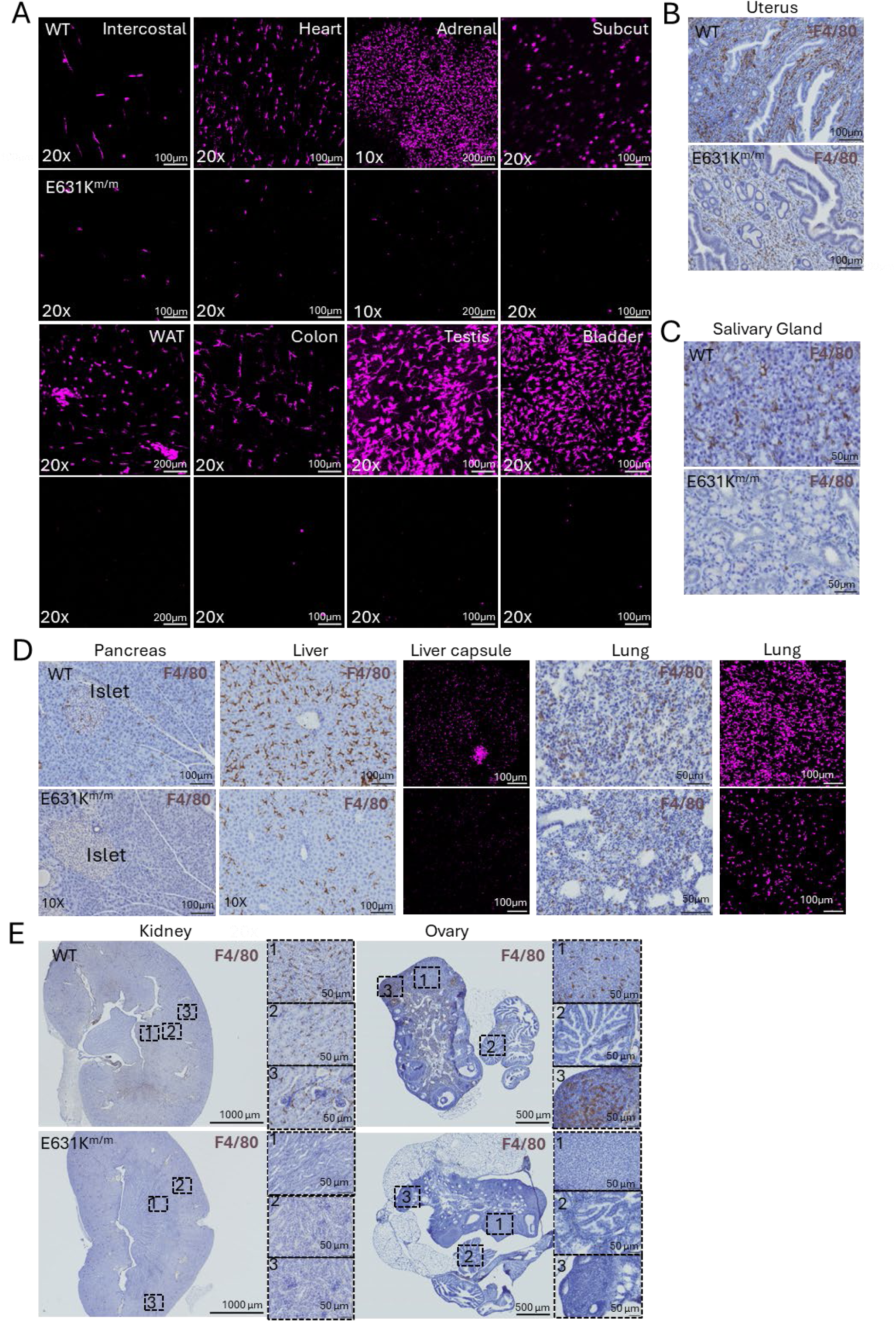
Analysis of tissue resident macrophages in WT and E631K^m/m^ mice. Panel A shows representative maximum intensity projections of *Csf1r-*EGFP signal (pseudocolored purple) in WT and E631K^m/m^ mice (intercostal= skeletal muscle; Subcut= subcutaneous fat; WAT= white adipose tissue; male epididymal fat pad). Panels B-E show representative images of F4/80 staining of tissues indicated. Inset panels in (E) show the loss of F4/80^+^ cells throughout the cortex and medulla in E631K^m/m^ kidney and in the ovary. Inset panels show the loss of macrophages throughout the ovary and fallopian tube in the E631K^m/m^ mice.

**Figure S2.**
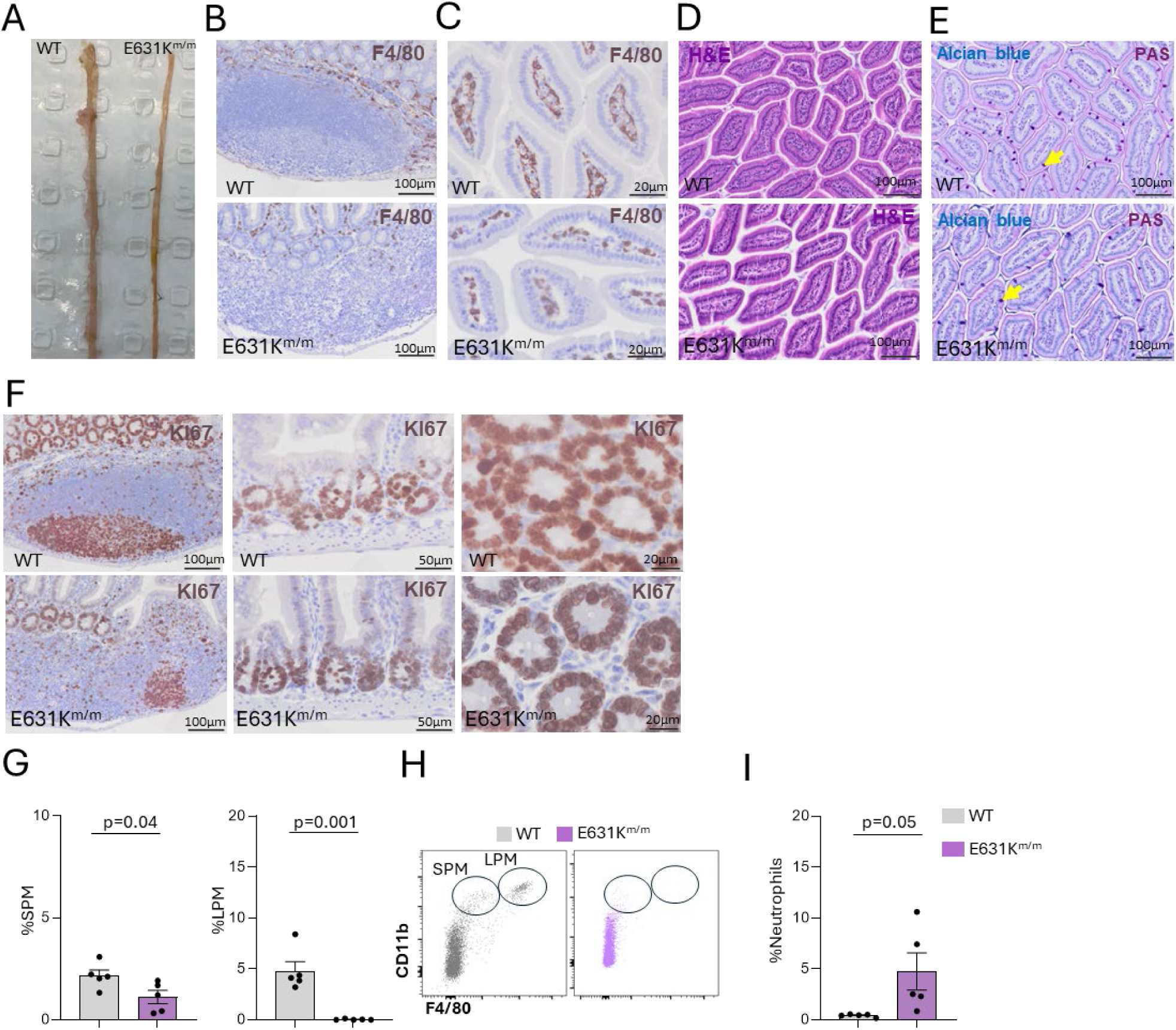
Comparative analysis of intestinal mucosa in WT and E631K^m/m^ mice. (A) Representative macroscopic images of small intestine of adult WT and E631K^m/m^ mice. (B-F) Representative images of intestinal sections stained for the markers indicated. PAS: Periodic Acid Schiff Stain. Yellow arrows indicate goblet cells stained with alcian blue. (G-I) Peritoneal cells were stained for CD11b, F4/80 and Ly6G and analysed by flow cytometry. Data show means and standard deviation. Each point is an individual animal. *p* values were determined by unpaired two-tailed t-test.

### Brain development in E631K^m/m^ mice is not affected by the absence of microglia

The brains of E631K^m/m^ mice lacked microglia detectable by staining for the IBA1 marker (**Figure 3A**) or the CD45^low^/CD11b^high^ population identified by flow cytometry of disaggregated tissue (**Figure 3B**). F4/80 staining in multiple brain regions confirmed the absence of both microglia and border-associated macrophage populations (**Figure 3C-E)**. Hence, the residual CD45^high^/CD11b^low^ cells detected in **Figure 3B** and F4/80^+^ cells associated with meningeal vessels (**Figure 3D**) likely includes monocytes. In E631K^m/m^ mice we noted accumulation of F4/80^+^ monocyte-like cells in the velum interpositum (**Figure 3F**), an extension of the meninges underlying the hippocampus implicated as a portal of myeloid cell entry to the brain ^30^. Despite the absence of microglia, there was no evidence of dysgenesis of the corpus callosum, involution of the olfactory bulb or ventricular enlargement reported in inbred *Csf1rko* mice ^31^ or any other gross anatomical change (**Figure 3C**). Consistent with analysis of *Csf1rko* mouse brain ^31^, localisation of GFAP revealed an increase in % area and fluorescence intensity, indicating a mild astrocytosis, mainly in the hippocampus, in E631K^m/m^ mice (**Figure 3G).**

**Figure 3.**
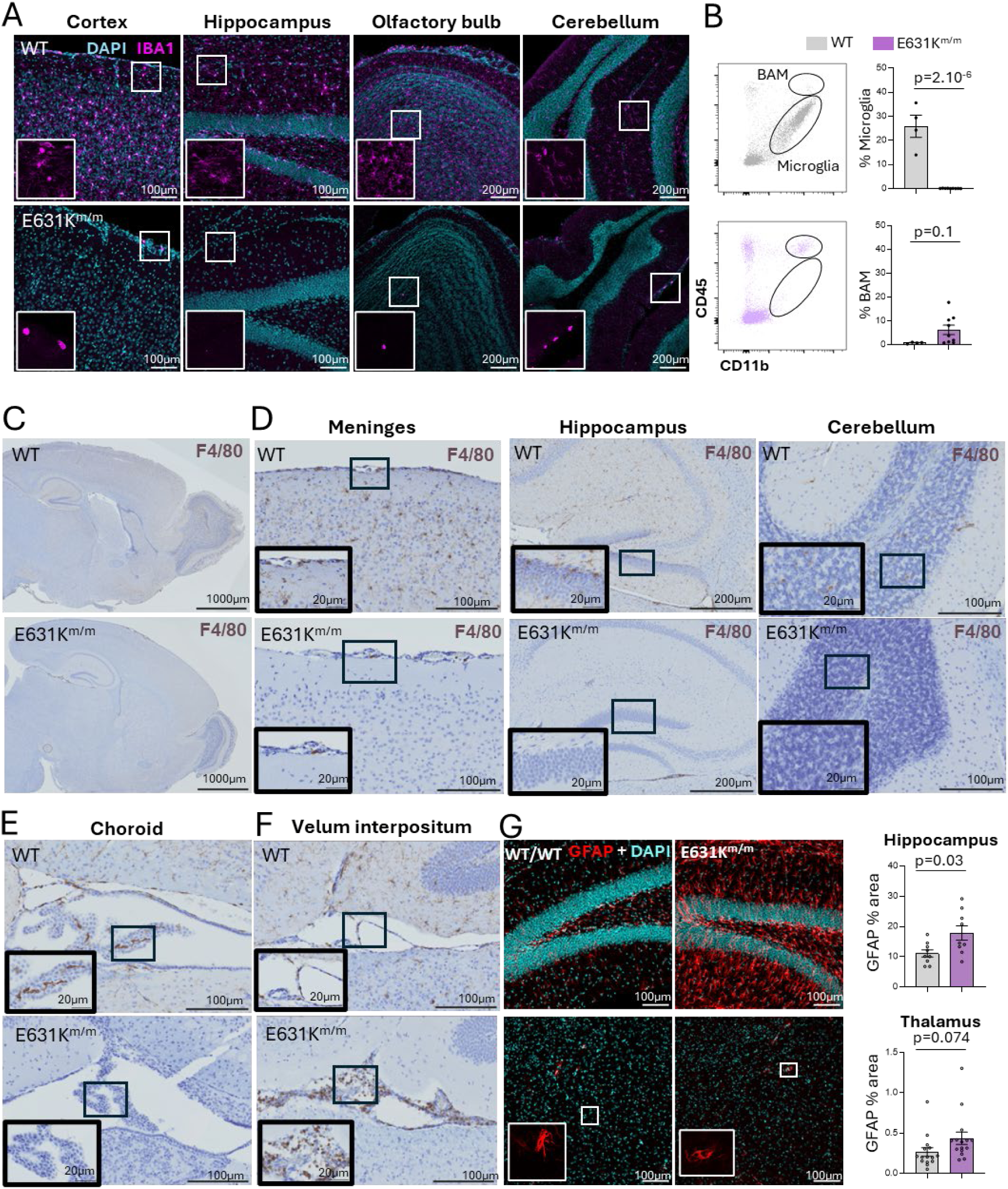
The effect of microglial deficiency in E631K^m/m^ mice. (A) Free floating brain sections from adult WT and E631K^m/m^ mice were stained for IBA1 and imaged by confocal microscopy. (B) Disaggregated brain cells were stained for CD45 and CD11b and analysed by flow cytometry. (C-F) Representative sections of selected brain regions from adult WT and E631K^m/m^ mice were stained for F4/80 and counterstained with hematoxylin. (G). Free floating brain sections from adult WT and E631K^m/m^ mice were stained for GFAP and imaged by confocal microscopy. Panels show representative images of hippocampus (upper) and thalamus (lower), quantified using Imaris (N=8). Each point is an individual animal. *p* values were determined by unpaired two-tailed t-test

### E631K^m/m^ mice are partly deficient in circulating monocytes

Automated hematology analysis showed that total peripheral white blood cell (WBC) count was not altered in E631K^m/m^ mice; but the mutants exhibited an absolute and proportional increase in granulocytes and a decrease in lymphocytes, with no change in monocytes, platelets or erythrocytes (**Figure 4A, Figure S3A**). To accurately phenotype and quantify monocytes, since monocytes in mice cannot be separated from granulocytes by automated analysers^32^, we utilised flow cytometry. Ly6G^+^ neutrophils and NK1.1^+^ cells were excluded and monocytes were gated as CD11b^+^/*Csf1r*-EGFP^+^ (**Figure 4B-C, Figure S3B-C**). *Csf1r*-EGFP^+^ was included in the gating strategy to ensure exclusion of NK cells, which do not express the *Csf1r-* EGFP transgene ^24^ (**Figure S3B,C**). We confirmed the increase in Ly6G^+^ neutrophils as a proportion of WBC (**Figure 4D**). Within the CD11b^+^/Ly6G^-^ population there was an increase in the proportion of *Csf1r-*EGFP^-ve^ cells (**Figure 4E**) which we speculate represent immature myeloid cells, and a corresponding decrease in the proportion of monocytes defined by *Csf1r-* EGFP expression (**Figure 4E**). As a percentage of WBC, monocytes were reduced by 60-70%, but the ratio of Ly6C^high^/Ly6C^low^ monocytes was not significantly altered in E631K^m/m^ mice (**Figure 4E**). CD115 expression was greatly reduced in both monocyte populations in E631K^m/m^ mice mutant compared to WT (**Figure 4C**), which may be due, in part, to the high level of circulating CSF1 **Figure 1J**). Classical monocytes can be derived from either granulocyte-macrophage-progenitors (GMP) or monocyte-dendritic cell progenitors (MDP) in bone marrow, both of which express CSF1R ^33^. Monocytes derived from these distinct progenitors may be distinguished by expression of CD319 (MDP) or CD177 (GMP) ^34^. By contrast to the reported excess of GMP monocytes reported in C57BL/6J mice, in WT F2 mice CD319^+^ (MDP) were predominant, around 30% of classical monocytes. Detection of both markers was reduced in classical monocytes of E631K^m/m^ mice (**Figure 4F,G**).

**Figure 4.**
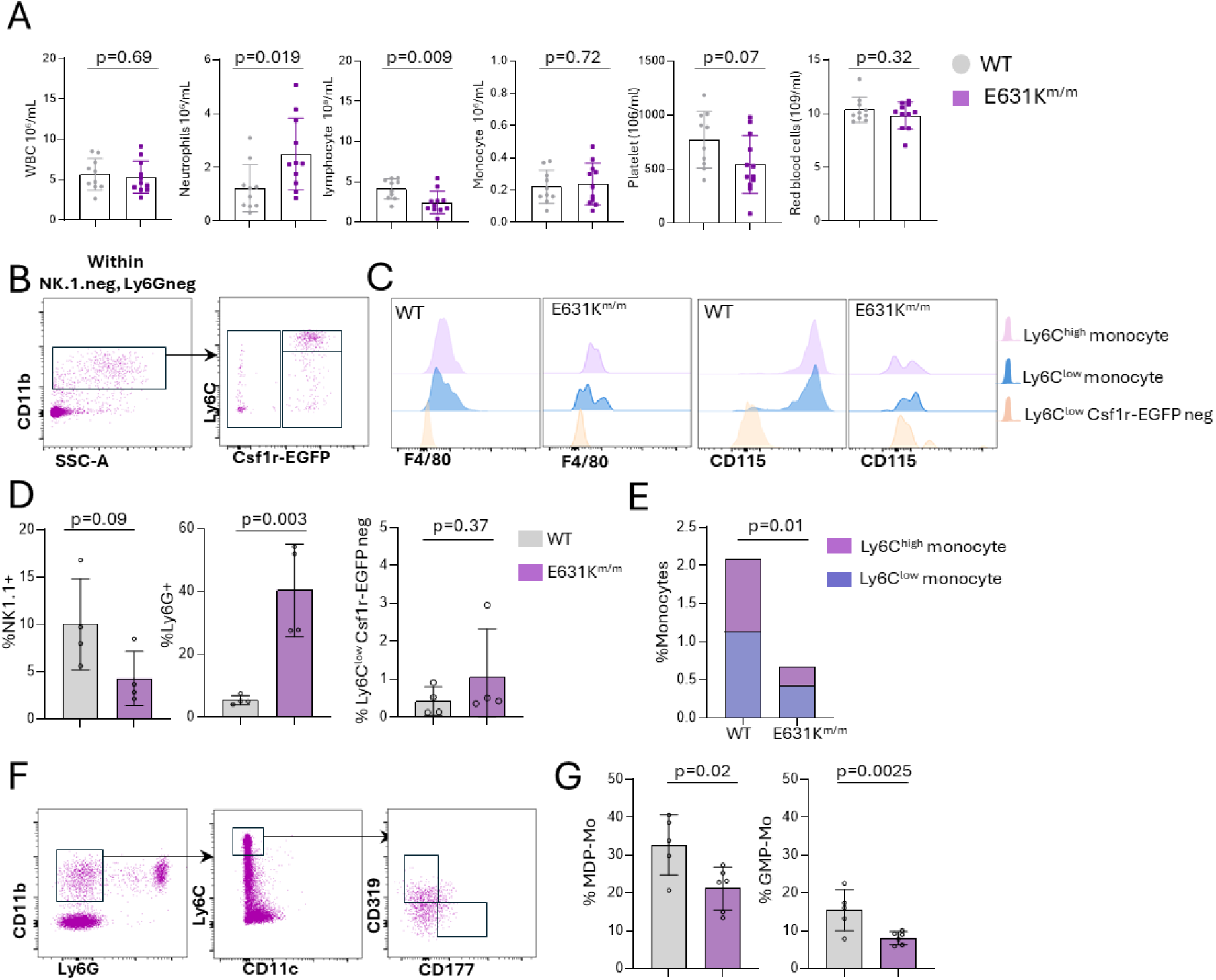
Analysis of blood cells in WT and E631K^m/m^ mice. (A) White blood cell (WBC), neutrophil, lymphocyte, monocyte, platelet and red blood cell, counts in peripheral blood were measured using the Mindray haematology analyser. (B) Representative flow cytometry gating strategy using *Csf1r*-EGFP to identify monocytes (gated on NK1.1^-ve^/Ly6G^-ve^ cells). (C) Representative profiles of F4/80 and CD115 expression on monocyte subsets and CD11b^+^/*Csf1r*-EGFP^-ve^ cells from WT and E631K^m/m^ mice. (D,E) Proportions of total white blood cells defined by surface markers indicated. In panel E, monocytes defined as NK1.1^-,^ Ly6G^-^, CD11b^+^, EGFP^+^ are separated as Ly6C^high^ or Ly6C^low^ as indicated. Data show means and standard deviation. Each point is an individual animal. *p* values were determined by unpaired two-tailed t-test

**Figure S3.**
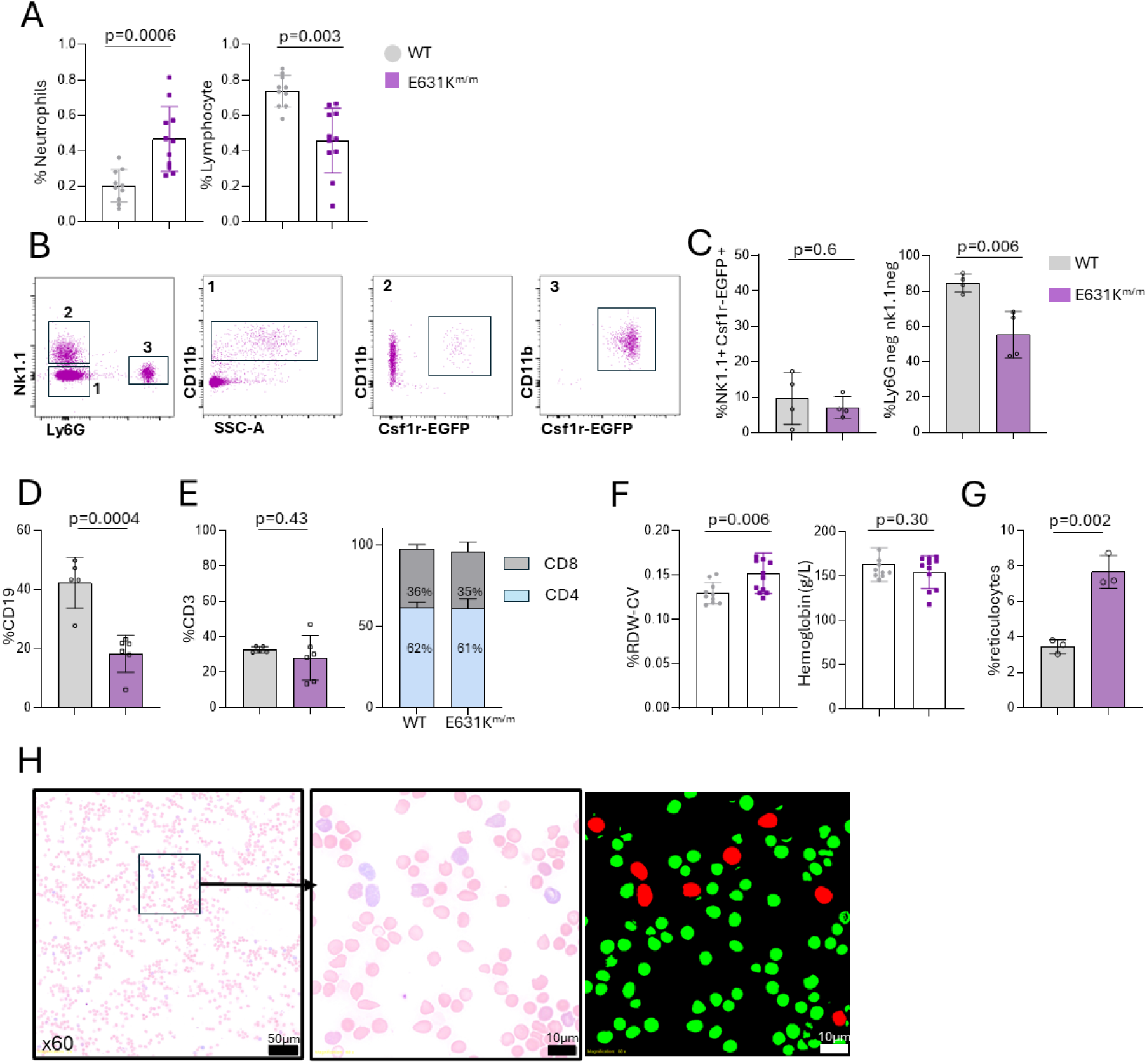
Analysis of blood leukocytes in WT and E631K^m/m^ mice (related to Figure 4). (A) Proportions of blood neutrophils and lymphocytes measured by haematology analysis expressed as a percentage of total cells. (B,C) Representative profiles of peripheral blood showing *Csf1r*-EGFP transgene expression in CD11b^+^ cells further defined as NK1.1^-^/Ly6G^-^ (region 1), NK1.1^+^ (region 2) and Ly6G^+^ (region 3) (D) Proportion of CD19^+^ B cells as a percentage of total blood leukocytes. (E) Proportions of CD4^+^ and CD8^+^ cells within the blood CD3+ T cell population. (F) Analysis of red cell distribution width-coefficient of variation (RDW-CV) and total hemoglobin. (G,H). Analysis of reticulocytes in peripheral blood. percentage of reticulocytes within blood smears (3 animals per genotype) stained with Wright Giemsa quantified by automatic segmentation. Data show means and standard deviation. Each point is an individual animal. *p* values were determined by unpaired two-tailed t-test.

The lymphocyte deficiency in E631K^m/m^ blood was specific to CD19^+^ B cells, whereas CD3^+^ T cell populations and CD4/CD8 ratio were unchanged (**Figure S3D-E**). Although red blood cell (RBC) count and hemoglobin were not affected, an increase in red cell distribution width (RDW-CV) and circulating reticulocytes in mutant mice (**Figure S3F-H)** suggests some dysregulation of RBC turnover.

### The impact of resident macrophage deficiency on bone marrow hematopoiesis

CD169 (SIGLEC1) is expressed by F4/80^+^ resident macrophages that form the centre of hematopoietic islands and osteal macrophages lining the bone surfaces ^33,35–38^. CD169^+^ cells were undetectable in E631K^m/m^ bone marrow. However, F4/80^+^ staining indicated retention of a subpopulation of macrophages (**Figure 5A).** Although resident macrophages cannot be assessed by flow cytometry due to the fragmentation that occurs with tissue disaggregation ^39^, we analysed bone marrow monocytes, adopting the same flow cytometry gating strategy as in peripheral blood (**Figure 5B)**. In WT mice, Ly6C^high^ monocytes and a proportion of Ly6C^low^ monocytes were CD115^+ve^, but CD115 was barely detectable in E631K^m/m^ marrow monocytes (**Figure 5C**). As observed in blood, Ly6G^+^ neutrophils and Cd11b^+^/*Csf1r-*EGFP^-^ putative immature myeloid cells formed a significantly greater proportion of the total cell population in E631K^m/m^ marrow but monocytes were not changed (**Figure 5D,E).** The mutant marrow contained relatively fewer CD19^+^ B cells, whereas the minor populations of T cells and NK cells were not altered (**Figure 5F-G**).

**Figure 5.**
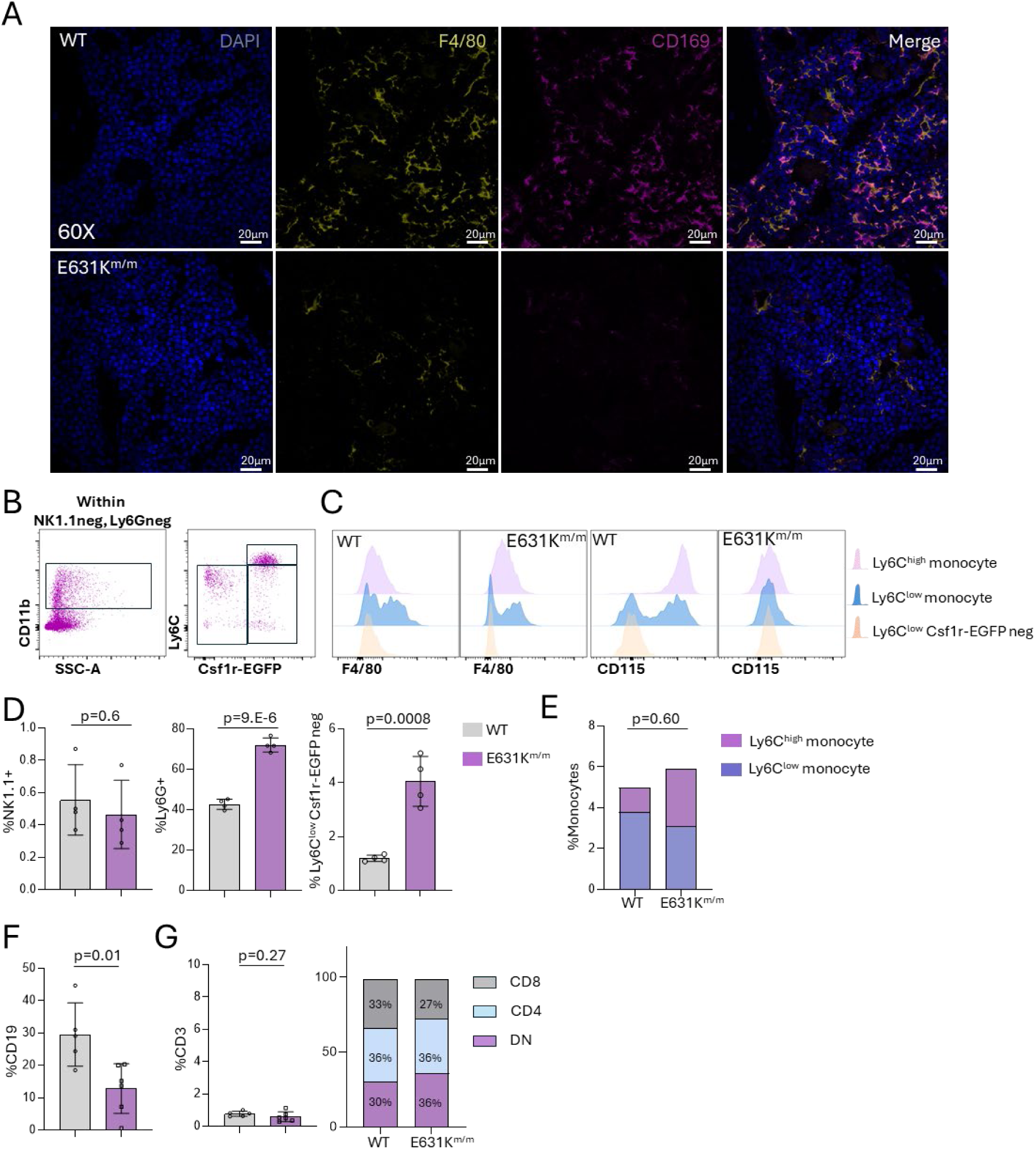
The effect of Csf1r mutation on bone marrow. (A) Sections of decalcified femurs from WT and E631K^m/m^ mice were stained for F4/80, CD169 and DAPI and imaged by confocal microscopy. Representative images are shown. (B) Representative flow cytometry gating strategy using *Csf1r-*EGFP for detection of bone marrow monocytes (gated on NK1.1^-ve^/Ly6G^-ve^ cells). (C) Representative profiles of F4/80 and CD115 expression on monocyte subsets and CD11b^+^/*Csf1r*-EGFP^-ve^ cells from WT and E631K^m/m^ mice. (D) Proportion of total bone marrow cells defined by the surface markers indicated. (E) Proportions of CD19^+^ B cells and (F) CD3^+^ T cells (subdivided based upon CD4 and CD8) in bone marrow measured by flow cytometry. Data show means and standard deviation. Each point is an individual animal. *p* values were determined by unpaired two-tailed t-test.

The increase in myeloid cell contribution to total bone marrow cellularity in mutant mice was reflected in a decreased proportion of lineage negative (Lin^-^) cells (**Figure 6A-F**). However, within the Lin^-^ population, KIT^+^ committed progenitors (LK) and KIT^+^/SCA1^+^ (LSK) hematopoietic stem cells (HSC) were increased (**Figure 6A-B)**. Within the LSK fraction, the proportion of long-term hematopoietic stem cells (LT-HSC, CD150^+^/FLT3^-^/CD48^-^) was reduced, while committed progenitor subpopulations, ST-HSCs and multipotent progenitors (MPPs) were unchanged (**Figure 6C-D)**. Wright Giemsa staining of bone marrow cells confirmed the marked accumulation of myeloblasts and granulocytic cells in E631K^m/m^ mice (**Figure 6E).** Cell cycle analysis showed a small reduction in S and G2/M phase cells in Lin-Kit+ committed progenitors in mutant mice (**Figure 6F**). Within the erythroid lineage progressive differentiation was assessed based on expression of TER119 and CD71 ^40^. Relative to WT, E631K^m/m^ marrow contained a reduced proportion of erythroid progenitors (R2, TER119^+^/CD71^+^) and increased erythroblasts (R4, TER119^+^/CD71^-^) (**Figure 6G, H**). To extend these observations we localised F4/80 and CD71 in sections of juvenile bone marrow. Consistent with the flow cytometry data the apparent density of CD71^+^ erythroblasts was reduced in E631K^m/m^ marrow. However, the remaining CD71^+^ cells appear associated with the residual F4/80^+^ macrophage population suggesting that erythropoiesis continues to be associated with erythroblastic islands despite the loss of CD169 expression (**Figure 6I, J**).

**Figure 6.**
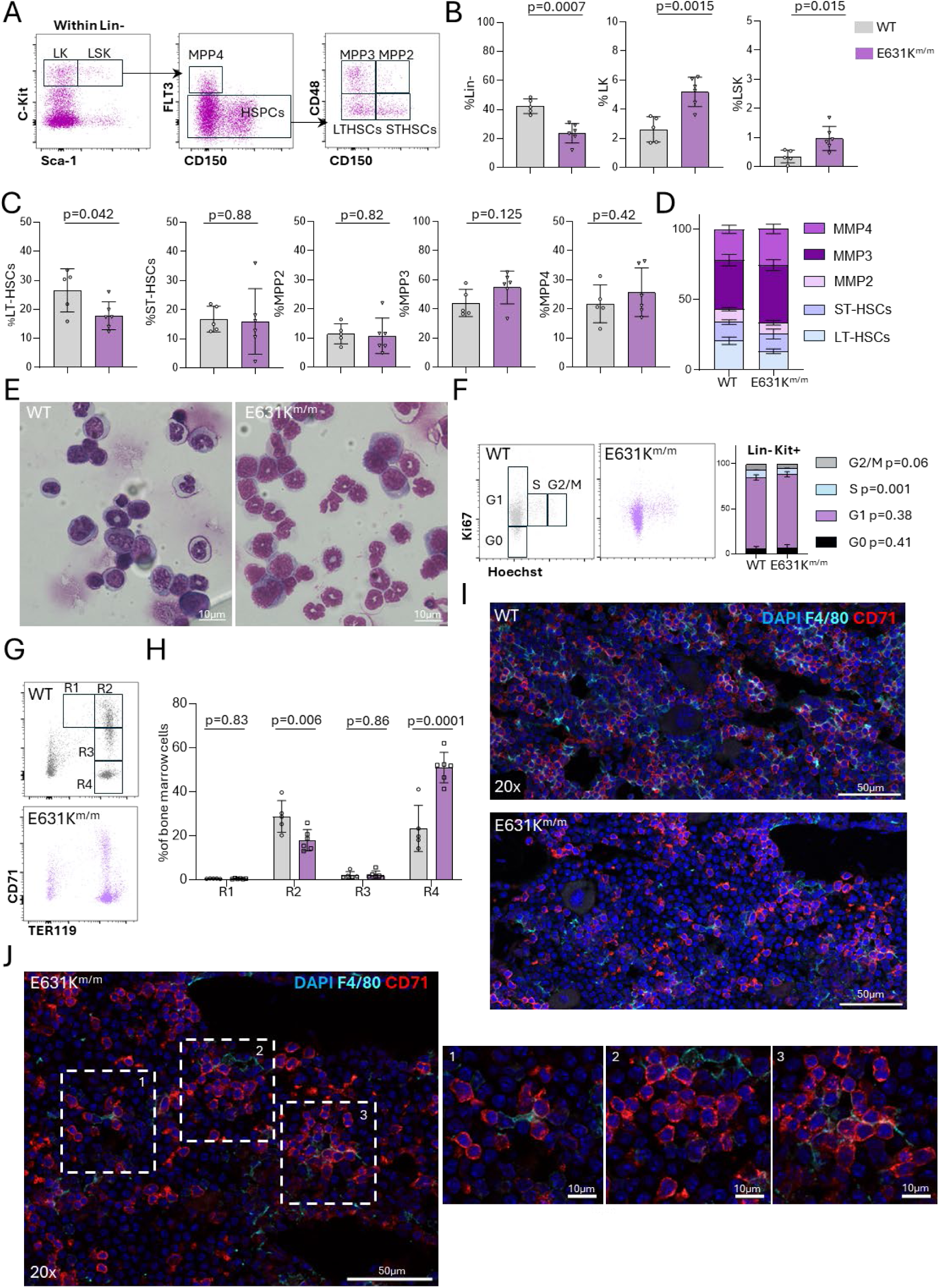
Hematopoietic stem and progenitor cell in bone marrow of E631K^m/m^ mice. (A) Representative flow cytometry gating strategy for bone marrow hematopoietic stem and progenitor cells. Lineage negative cells were identified as Kit^+^ (LK, committed progenitors) and Kit^+^/Sca1^+^ (LSK, hematopoietic progenitor cells). LSK were further divided based upon detection of FLT3, CD48 and CD150 as indicated. (B) Graphs show the proportion of Lin-, LK and LSK as a percentage of the total bone marrow. (C,D) Proportions of LT-HSC, ST-HSC, MPP2, MPP3 and MPP4 within the LSK population. (E) Representative images of bone marrow cytospins from WT and E631K^m/m^ mice stained with hematoxylin and eosin. (F) Cell cycle analysis in marrow progenitor cells of WT and E631K^m/m^ mice Representative flow cytometry gating strategy and quantification of LK cells in G0, G1, S and G2/M phases of the cell cycle based on Hoechst 33342 staining. (G, H). Analysis of erythroid differentiation based on CD71 and Ter119 staining. (I-J) Colocalisation of CD71^+^ erythroblasts and F4/80^+^ macrophages in juvenile (3 week) bone marrow. Data show means and standard deviation. Each point is an individual animal. *p* values were determined by unpaired two-tailed t-test.

### Extramedullary hematopoiesis in the spleen

The gross architecture of the spleen appeared unaffected in E631K^m/m^ mice and F4/80^+^ red pulp macrophages were retained (**Figure 7A-B**). However, the marginal zone was profoundly disrupted in E631K^m/m^ mice, marked by the complete loss of CD169^+^ metallophilic macrophages and reduction and disorganisation in the CD209b^+^ (SIGNR1) marginal zone macrophage population ^41^ (**Figure 7C**). We analysed spleen populations using the same flow cytometry gating strategy as in blood and marrow (**Figure 7D, Figure S4A-B)**. CD115 was barely detectable in freshly isolated E631K^m/m^ splenic monocytes (**Figure 7E**) but was increased in following incubation *in vitro* to allow dissociation of bound ligand and re-expression (**Figure S4C**). The Ly6C^low/-^ population in spleen was predominantly F4/80^high^ in WT mice (**Figure 7E**) and likely includes red pulp macrophages which appeared reduced in the mutant. However, as noted with respect to the bone marrow analysis, recovery of resident macrophages following tissue disaggregation is not quantitative. As observed in blood and marrow, there was an increase in Ly6G^+^ neutrophils and putative immature myeloid cells (Ly6G^-^/NK1.1-/CD11b^+^) in E631K^m/m^ spleen (**Figure 7F**). Both Ly6C^high^ and Ly6C^low^ monocytes were also proportionally increased (**Figure 7F**). Consistent with a previous report ^12^, classical DC (CD11c^+^/MHCII^high^) were not affected by *Csf1r* mutation (**Figure S4D**). As in blood and bone marrow, splenic CD19^+^ B cells were significantly reduced and T cells were unchanged in E631K^m/m^ mice (**Figure S4 E-F**).

**Figure 7.**
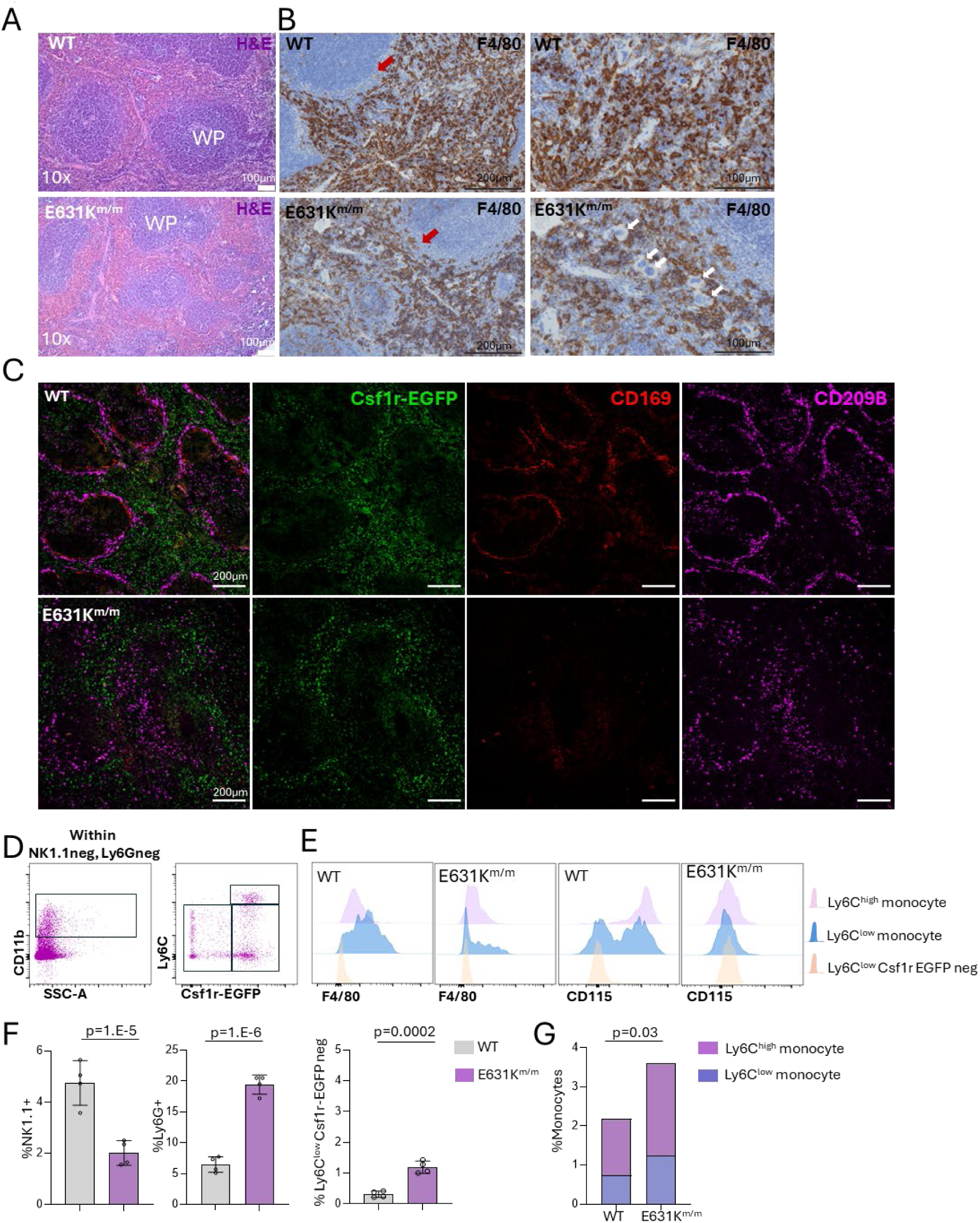
Splenic leukocyte populations in E631K^m/m^ mice. (A) Hematoxylin and eosin (H&E) staining and (B) Immunolocalisation of F4/80 in WT and E631K^m/m^ adult spleen. Representative images are shown. Red arrows: marginal zone. White arrows: megakaryocytes. (C) Representative images of immunolocalisation of marginal zone metallophils (CD169), marginal zone macrophages (CD209b) and *Csf1r-*EGFP in WT and E631K^m/m^ adult spleen. (D) Representative flow cytometry gating strategy for splenic monocytes. (E) Representative profiles of F4/80 and CD115 expression on monocyte subsets and *Csf1r*-EGFP^-ve^ cells. (F) Proportion of total spleen cells defined by surface markers indicated. Data show means and standard deviation. Each point is an individual animal. *p* values were determined by unpaired two-tailed t-test.

**Figure S4.**
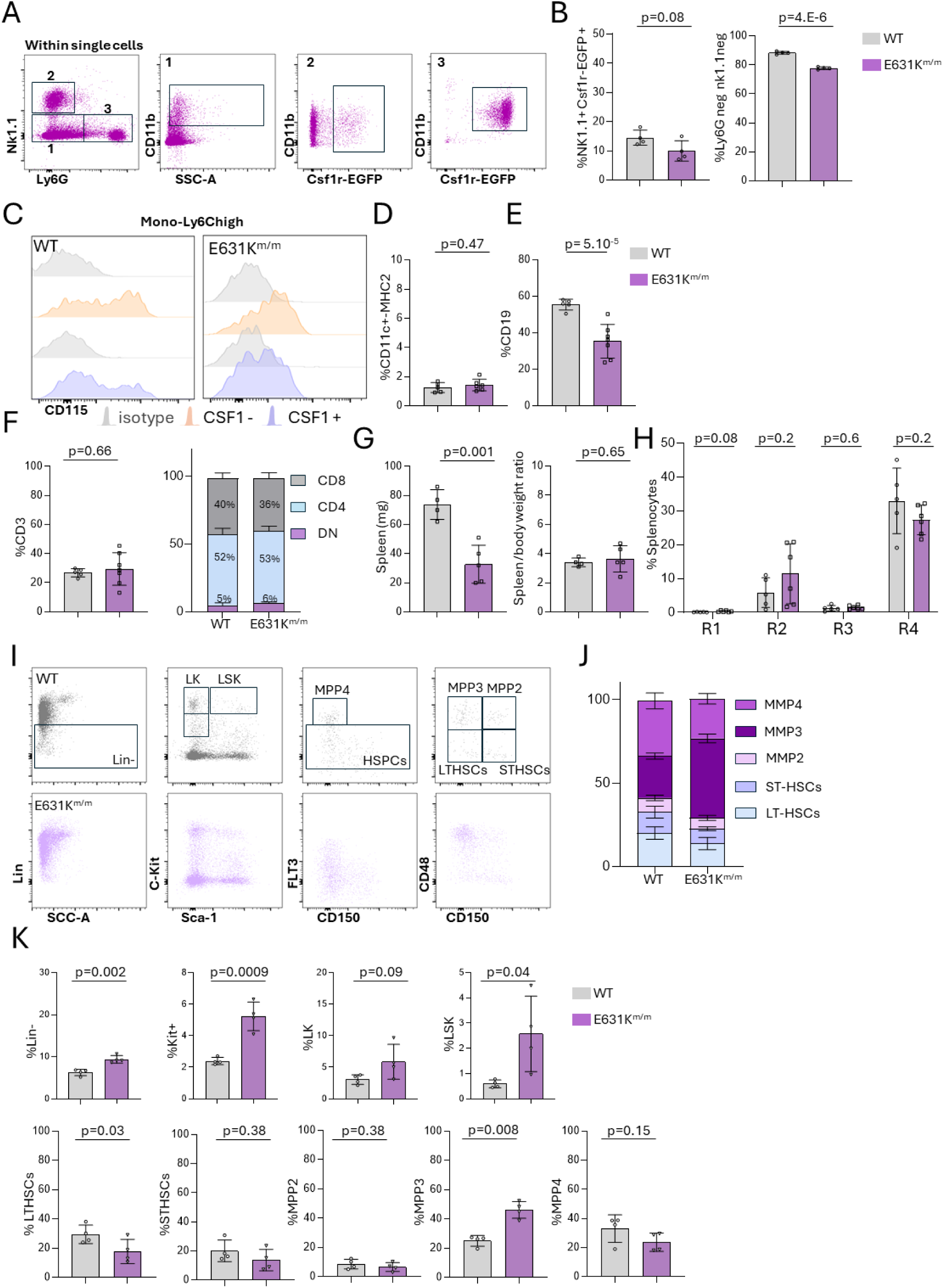
Analysis of splenic leukocytes in WT and E631K^m/m^ mice (related to Figure 7). (A, B) Representative flow cytometry plots showing the expression of the *Csf1r*-EGFP reporter on defined populations in spleen (gated on live, single cells). (C) Freshly isolated splenocytes were washed and incubated at 4°C for 90 min to facilitate CD115 surface expression, then stimulated with 100 ng/ml CSF1 for 1 hour at 37°C followed by staining for myeloid markers, including CD115 (D-F) Percentages of dendritic cells (CD11c^high^/MHC2^high^), CD19^+^ B cells, CD4^+^ and CD8^+^ T cells as a percentage of total spleen cells. G) Absolution spleen weight and spleen/body weight ratio in adult WT and E631K^m/m^ mice. (H) Analysis of erythroid progenitor populations in spleen (gating strategy in Figure 6G). (I-K) Analysis of progenitor populations in the spleen of WT and E631K^m/m^ mice. Data show means and standard deviation. Each point is an individual animal. *p* values were determined by unpaired two-tailed t-test.

Megakaryocytes, a hallmark of extramedullary haematopoiesis (EMH), were readily detected in the splenic red pulp of E631K^m/m^ mice (**Figure 7B**). Splenic EMH was reported in *Csf1rko* mice, marked by an increase in high proliferative potential colony forming cells (responsive to IL6, Stem Cell Factor, IL3 and CSF2) and erythroid burst-forming units, associated with splenomegaly ^10^. In contrast to *Csf1rko* mice, the spleens of E631K^m/m^ mice were reduced in size, proportional to their lower body weight (**Figure S4G)**. We found no change in the distribution of erythroid progenitor cells **(Figure S4H)**, but identified accumulation of Lin^-^, Lin^-^Kit^+^, and HSPCS (LSK and LK) in E631K^m/m^ spleens (**Figure S4I**). Among LSKs, we observed a reduced proportion of LT-HSC and an increase in MPP3, myeloid-biased progenitors that are the main contributor to myelopoiesis ^42,43^ (**Figure S4J-K**).

## Discussion

### E631K^m/m^ is a null mutation with variable penetrance dependent on genetic background

Because of the perinatal lethality of the *Csf1rko* on the most widely-used genetic background, C57Bl/6J ^12,44^, there have been few analyses of the impact of congenital CSF1R deficiency in mice since the original study in outbred mice ^10^. The global loss of macrophages and osteoclasts and functional analysis in **Figure 1**, support previous evidence that *Csf1r*^E631K^ is a loss of function mutation ^20,21,23,45^. Hence, differences in phenotype of homozygotes compared to previous studies of *Csf1rko* mice can be attributed to genetic background. We have demonstrated this directly by mitigating the prenatal lethality of the mutation on the C57BL/6J background by crossing to BALB/c. As noted in the introduction, the microglia-deficient *Csf1r*-FIRE mutant mouse is also long-lived and fertile on a mixed genetic background ^46^ but the same mutation causes perinatal loss and hydrocephalus on the C57BL/6J background ^47^. These findings add further weight to the view that C57BL/6J mice may not provide a generalisable model of macrophage biology ^48^.

### The impact of macrophage deficiency on mouse postnatal development

Analysis in Figure 2 and S2 confirmed the complete depletion of macrophages in most organs of the E631K^m/m^ mice. As in the rat *Csf1rko*^26^, circulating C1q provided a sensitive biomarker of the macrophage deficiency, reduced to the limits of detection in mutant mice. Previous studies in mouse inferred that macrophages were the major source of C1q ^49^.

Macrophages are proposed to regulate development and homeostasis in every major organ system ^50–52^. In a comprehensive analysis of gene expression in tissues of juvenile *Csf1rko* rats we found evidence of postnatal developmental delay likely due in part to complex interactions between bone, liver and pituitary leading to systemic changes in circulating regulators ^26^. In both rats and mice there is partial retention of macrophages in the bone marrow, lung, liver, intestine and spleen. We confirmed in mouse the complete loss of CD169^+^ marginal zone metallophils in the spleen reported in the *Csf1rko* rat ^26^. These cells are absent in *Csf1*^op/op^ mice ^53^ and even heterozygous *Csf1r*^E631K^ mutation is sufficient to deplete them ^23^. By contrast, the CD209B^+^ (SIGNR1) marginal zone macrophages ^54^, which were CSF1R-dependent in the rat ^26^ were retained but disorganised in the mutant mice. The species-specificity may relate to the substantial difference in splenic architecture between mice and rats ^55^. Given their intimate association with sinus endothelium, we speculate that the disrupted distribution of CD209B^+^ macrophages in mutant mice reflects underlying vascular changes. In both species, red pulp macrophages are partly CSF1R-independent.

There are several other differences between CSF1R loss of function mutations in rats and mice. *Csf1rko* mice on an inbred FVB/J background, where pups are born but few survive to weaning, have severe neurodevelopmental impacts including callosal agenesis, hydrocephalus and involution of the olfactory bulb ^31,56,57^. Similar neurodevelopmental impacts were also observed, albeit later onset, in the *Csf1rko* rat ^58^ and in human patients with homozygous *CSF1R* mutations ^8,9^. By contrast, the only impact of the mouse E631K^m/m^ mutation in the brain on an F2 background was a mild astrocytosis. It remains to be seen whether these mice would develop the age-dependent neuropathology, including calcification of the thalamus, seen in microglia-deficient mice ^59,60^.

Developmental delay in the *Csf1rko* rat liver is associated with steatosis whilst visceral adipose tissue is almost completely absent ^5,26^. Neither phenotype is evident in E631K^m/m^ mice (**Figure 2A,D**). The kidneys of E631K^m/m^ mice are histologically normal (**Figure 2E**) by contrast to the selective dysplasia of the medulla observed in *Csf1rko* rats ^4^. Despite retention of bronchoalveolar (but not interstitial) lung macrophages, *Csf1rko* rats succumb to an emphysema-like lung pathology ^26^ which was not seen in the E631K^m/m^ mice (**Figure S2D**). Some of the species differences may be related to regulation of IGF1, which was almost undetectable in *Csf1rko* rats ^4,26^ but reduced by only 50% in E631K^m/m^ mice.

Previous studies suggested that *Csf1r* was expressed in intestinal epithelial cells and directly regulated Paneth cells and epithelial proliferation ^61–63^. Subsequent analysis indicated that CSF1R expression is macrophage-restricted in the gut ^33^ but depletion of macrophages using anti-CSF1R antibody nevertheless altered epithelial differentiation ^64^. By contrast, we did not detect any overt perturbation in intestinal villus histology in E631K^m/m^ mice. Mucosal thickness was greatly reduced, which is difficult to interpret given the size differences between WT and mutant mice. E631K^m/m^ mice retained monocyte-like cells in the submucosa and lamina propria which may continuously replace intestinal macrophages ^65^, likely enabling the maintenance of intestinal homeostasis. Expression profiling of intestine in the *Csf1rko* rat was consistent with a reduction in mature macrophages but also did not reveal any change in the relative abundance of any epithelial population ^26^.

### The contribution of resident bone marrow macrophages to the regulation of hematopoiesis

The resident macrophages of bone marrow and foetal liver interact intimately with stem cells and proliferating progenitors and have long been considered an essential component of the haematopoietic niche ^37,38,66–70^. There have also been reports that CSF1R is expressed by hematopoietic stem cells ^67,71,72^ although this is likely to be an artefact of tissue disaggregation ^73^ and is contradicted by analysis of expression of a knock-in CSF1R reporter transgene ^33^. Deletion of CD169^+^ erythroblastic island macrophages using a CD169-DTR transgene was reported to lead to selective loss of erythroblasts ^74^. Romano *et al.* ^70^ provided evidence that erythroblastic islands also contain myeloblasts and suggested that the island-associated macrophages regulate the balance between erythropoiesis and granulopoiesis, similar to the reported function of CSF1R-dependent macrophages in the HSC niche in fetal liver ^69^. The colocalisation of F4/80 and CD71 in E631K^m/m^ marrow indicates that erythropoiesis remains at least partly associated with residual erythroblastic island (EBI) macrophages. The loss of detectable CD169 in the marrow suggests that the phenotype and likely function of these residual macrophages is altered by the lack of CSF1R signalling, and that CD169 is not required for erythropoiesis. Interestingly, CSF1 treatment of mice causes neutrophilia alongside the monocytosis ^75,76^. This response might be mediated indirectly through effects on the EBI macrophages. Further analysis of the phenotype of CSF1R-independent bone marrow macrophages is constrained by their extensive ramification and the difficulty of isolating intact macrophages from marrow ^39^.

Previous studies based on antibody blockade of CSF1R or CSF1^76–79^, CRE-mediated conditional deletion of *Csf1* in the marrow or vasculature ^80–82^ or inducible *Csf1r* deletion ^83^ concluded that non-classical (Ly6c^low^) monocytes are selectively dependent upon CSF1R signalling. Circulating monocytes were reduced in E631K^m/m^ mice, as reported in *Csf1rko* mice ^10^ and rats ^4^, but the Ly6C^high/low^ ratio was not significantly affected. Based upon the effect of anti-CSF1R antibody, CSF1R signalling is not required for monocytopoiesis ^78,84^ and monocytes were not proportionally depleted in E631K^m/m^ bone marrow or spleen. FLT3L might contribute to the maintenance of monocytopoiesis in the absence of CSF1R signals. *FLT3L* mutation is associated with monocyte deficiency in humans but not in mice ^85,86^. Like CSF1^76^, FLT3L induces a profound monocytosis when administered to mice ^87^ and has been shown to support osteoclast development in *Csf1*^op/op^ mice ^88^. On the other hand, monocyte release from the bone marrow ^89^ and circulating monocyte half-life and extravasation^90,91^ are regulated processes that may be affected by tissue macrophage depletion contributing to the monocytopenia observed.

Although CSF1R is expressed by committed myeloid progenitors (CMP, GMP) ^33^, the granulocytosis observed in E631K^m/m^ and *Csf1rko* mice is not likely to involve a direct impact on progenitor fate. In *Csf1rko* rats, granulocytosis was reversed by WT bone marrow cell transfer without any contribution of donor cells to this lineage ^4,6^. Mature granulocytes are retained in the marrow for 4-6 days but have a half-life of hours in the circulation. The macrophages of the bone marrow are the major site of granulocyte clearance compared to liver and spleen ^92,93^. Hence, the increase in mature granulocytes in the marrow, blood, peritoneum and spleen of macrophage-deficient mice could also be a function of reduced release or clearance. The same argument may be applied to the erythroid lineage with reduced clearance of senescent red cells by liver and splenic macrophages ^94^ leading to increased circulating half-life, increased red cell distribution width (RDW-CV) and feedback regulation of erythropoiesis. The selective loss of B lymphocytes in marrow, blood and spleen in mutant mice was also evident in *Csf1rko* mice and rats ^1,4,5,10^. Although *Csf1r* may be expressed in a B cell progenitor in fetal liver ^95^, the loss of B cells is also unlikely to be cell-autonomous. Wild-type bone marrow transfer in *Csf1rko* rats also corrects the B cell deficiency without donor contribution to the B cell lineage ^6^. The lymphopoietic niche in bone marrow is dependent upon osteoblasts (reviewed in ^66^). Hence, the disruption of B lymphopoiesis associated with macrophage deficiency may reflect their role in the regulation of osteoblast function ^35^.

Although the relative proportions of stem and progenitor cells in the bone marrow were not significantly altered in the E631K^m/m^ mice, the absolute cellularity is clearly decreased due to osteopetrosis. The original analysis of CSF1R-deficient mice noted evidence of extramedullary hematopoiesis in the spleen, probably a consequence of marrow insufficiency ^10^. The more detailed analysis of the progenitor populations in the spleen of E631K^m/m^ mice confirmed this observation. However, unlike *Csf1r^-/-^* mice ^10^, the E631K^m/m^ mice did not have enlarged spleens so the overall contribution of the spleen to blood homeostasis is uncertain.

### Defined crossbred mouse lines as a model for the study of macrophage biology

We have demonstrated that perinatal lethality and hydrocephalus associated with *Csf1r* mutation depend upon inbred genetic background. The effect of genetic background is not due to strain-specific effects on macrophage survival; our analysis demonstrates almost complete loss of resident tissue macrophages in most organs of the F2 E631K^m/m^ mice. The likely explanation is that F2 mice are heterozygous for many of the thousands of coding and non-coding variants that distinguish the parent strains and genetic diversity provides increased resilience ^96,97^. Inbred lines are commonly used in research to minimise inter-individual phenotypic variation but this predicted outcome is not actually evident in published data ^98^. More importantly, many genetic resources have been generated on the C57BL/6J background. F2 mice provide a way to use those resources whilst mitigating the poor resilience of inbred mice. In our study we chose BALB/c as the breeding partner. Differences in macrophage function and gene expression between the two strains have been widely recognised ^48,99,100^. As exemplified by the *Csf1r* mutation, any allele of interest generated in C57BL/6J mice can be introgressed into an F2 background. The homozygous *Csf1r* mutation may also be used to generate macrophage-specific chimeras. In the rat and in *Csf1r*-FIRE mutant mice, intraperitoneal transfer of WT bone marrow cells at weaning allowed complete repopulation of tissue macrophages without contribution to the blood monocyte pool ^26,101^. MHC genotyping is straightforward ^102^; depending upon their genotype, F2 mice can receive MHC identical bone marrow from F1 or parent strains. In summary, we suggest that defined F2 strains offer a tractable and more informative alternative to inbred mice for the study of macrophage biology.

### Conclusion

We have presented a comprehensive analysis of the impact of homozygous *Csf1r* mutation in mice. Whilst osteopetrosis and reduced somatic growth are conserved features of CSF1R mutations in outbred mice, rats and humans^2^, in outbred mice many developmental and homeostatic functions of macrophages appear unaffected when these cells are absent.

## STAR Methods

### Animal breeding

In the original generation of the *Csf1r*^E631K/+^ line (C57BL/6J.Csf1r^Em1Uman^ (Tg16)) donor and recipient mice were C57BL/6JOlaHsd. The progeny backcrossed to C57BL/6JCrl were transferred from the UK, rederived, bred and maintained in specific pathogen free facilities at the University of Queensland (UQ) facility within the Translational Research Institute. To enable visualisation of myeloid populations in tissues, the *Csf1r*^E631K^ line was bred to the *Csf1r*-EGFP reporter transgenic line ^24^ backcrossed >10 times to the C57BL/6JArc genetic background. Genotyping of the mouse lines was carried out as described previously ^23^. BALB/c mice were obtained from the Animal Resource Centre, Perth, WA. All studies were approved by the Animal Ethics Committee of the University of Queensland. Mice were housed in individually ventilated cages with a 12 h light/dark cycle, and food and water available *ad libitum*.

### Blood analysis

Cell counts were performed on blood collected into EDTA-coated tubes (Greiner K3, Interpath, Australia) using a Mindray (BC-500) haematological analyser. Peripheral blood smears were stained with Wright-Giemsa (BIO Scientific, Australia) and scanned on an Olympus VS200 slide scanner with a 63x oil objective and automated oil dispersion utilising the VS200 ASW control software v3.3. Images were saved in .vsi format. A total of 16 200 x 200 µm regions of interest (ROIs) from blood smears were exported from the whole slide images in the VS200 Desktop software v3.3 in .tif format. The Olympus DNN module Detect was used to generate training labels to identify RBCs, reticulocytes and background pixels on the training images. The marked training images were saved and then loaded into the Olympus DNN learning module. An RGB specific network was run as the training configuration with a U-Net network architecture. 50,000 iterations were performed and the DNN was saved to apply to the .vsi raw data sets. The Neural Network file was saved and loaded back into the VS200 software. Neural Network Segmentation was then performed on random 1000 x 1000 µm ROIs from the .vsi images from the both the WT and E631K^m/m^ scans. The VS200 count and measure unit was used to measure and record the segmentation.

### Cell isolation and culture

Bone marrow was flushed from adult long bones by syringe with 2% fetal bovine serum (FBS) in phosphate buffered saline (PBS). Because of the osteopetrosis in the E631K^m/m^ mice the yield of cells was reduced by >50% compared to WT. Bone marrow cytospins were generated from 5×10^4^ cells, stained with Wright-Giemsa and imaged using a Nikon 50i microscope. To generate bone marrow-derived macrophages (BMM), ca. 10^7^ cells were seeded on 100 mm square bacteriological plastic (Sterilin, Thermo-Fisher, Australia) in 25 ml of complete medium (RPMI + 10% FBS, 25 U/mL penicillin, and 25 μg/mL streptomycin (Gibco, Thermo-Fisher, Australia) and differentiated for 7 days with the addition of recombinant CSF1-Fc^75^ (100 ng/ml) or recombinant mouse CSF2 (GM-CSF, R&D Systems, Minneapolis, Mn, USA; 50 ng/ml). For monitored differentiation assays 5×10^4^ bone marrow cells were seeded in 96 well coated plates in complete medium supplemented with 50 ng/ml mouse CSF2. Cell confluence was monitored every 6 h for 7 days using the Incucyte (Sartorius, Australia) cell-by-cell analysis system. Resazurin (Cayman Chem, Mi., USA; Cat #14322) was added at a final concentration of 0.025 mg/ml followed by incubation at 37°C/5% CO_2_ for 2-4 h before fluorescence was read (Ex: 530-540 nm, Em: 585-595 nm) using a POLARstar Omega plate reader (BMG Labtech). For osteoclast culture bone marrow cells were harvested as described above and seeded at 5 x 10^4^ cells/ml in 12 well tissue culture plates in complete medium with CSF1 (100 ng/ml) for 3 days. From day 3 the medium was replaced daily with medium containing CSF1-Fc (100 ng/ml) and RANKL (R&D Systems, 40 ng/ml) until day 7. Tartrate-resistant acid phosphatase (TRAP) staining was performed in situ using a kit (Cat #387A, Sigma, Australia) according to the manufacturer’s instructions. Peritoneal cells were obtained by flushing the peritoneal cavity with 10 ml PBS and prepared for flow cytometry staining by centrifugation at 400×g for 5 min at 4 °C prior to resuspension in FACS Buffer (FB, PBS containing 2% endotoxin-free FBS). Splenocytes were isolated by gently dissociating tissue with a 5 ml syringe in PBS, passed through a 40 µm filter, pelleted, subjected to red cell lysis (PharmLyse^TM^, BD Biosciences, Australia), washed and resuspended in FB. For brain cell isolation brains were dissected and minced with scissors in a 50 ml Falcon tube with 10 ml collagenase solution (HBSS, 20 µl 10 mg/ml DNase1 (Roche), 10 mg collagenase IV and 1 mg dispase (Life Technologies) and incubated in a shaking incubator at 37 °C for 45 min, 200 rpm. After incubation, brains were mashed through a 70 μm cell strainer with a syringe plunger into a 50 ml falcon tube and 25 ml FB was added. Cells were centrifuged at 400 g for 5 min at 4 °C. Supernatant was removed and the pellet resuspended in 12.5 ml isotonic Percoll (4.22 ml Percoll (GE Healthcare, Australia), 0.47 ml 10x PBS, 7.82 ml 1x PBS). Cells were centrifuged at 800g/4°C for 30 min with low acceleration and no brake. The myelin layer and supernatant was removed, and the pellet resuspended in 2 ml cold PharmaLyse^TM^ on ice for 10 min. 10 ml of FACS buffer was added and the cells centrifuged at 400g for 5 min at 4°C. The supernatant was removed, and the pellet resuspended in 1 ml of FB for flow cytometry staining. For cell cycle analysis, samples were incubated in Hoechst 33342 (BD, 5 μg/ml in PBS) for 5-10 mins then washed 3x in PBS.

### Flow cytometry

Following red cell lysis, single-cell suspensions from blood, bone marrow and splenocytes were stained with fluorophore-conjugated antibodies on ice and analysed on an LSR Fortessa, (BD, Australia). Cell-surface markers used to define hematopoietic cell types and antibodies are provided in Supplemental Table 1. Data were analysed using FlowJo (BD). Intracellular staining was performed using FIX & PERM kit (ThermoFisher, Australia).

### Whole mount imaging

Whole mount imaging of the *Csf1r*-EGFP reporter was performed using an FV3000 microscope (Olympus) and FLUOVIEW (Olympus) software. Tissue was harvested and kept in ice-cold PBS until imaging. Tissue was placed flat on a glass petri dish. Z-stack images were taken using the AF488 laser and combined to produce maximum intensity projections. For tissues with lower laser penetrance like heart, kidney and peritoneum, tissue was cut to obtain a representative surface. Images were taken using the same settings across genotype at 10x or 20x magnification and processed on ImageJ software v1.54.

### Immunofluorescence staining

For free-floating sections, brains were dissected from the skull and immersion fixed in 4% PFA for 5 hours at room temperature (RT), washed once in PBS and stored in PBS with 0.1% sodium azide. Brains were serially sectioned at 30μm in the sagittal plane using a Leica VT1200S vibratome. Sections were permeabilised (1% Triton-X100, 0.1% Tween-20 in PBS) for 1 h at RT, blocked (4% serum, 0.3% Triton-X100, 0.05% Tween-20 in PBS) for 1 h at RT, and then stained with primary antibodies (rabbit anti-IBA1 (1:500, Novachem, 019-19741; chicken anti-GFAP (1:500, Invitrogen, PA1-10004)) diluted in blocking buffer overnight at 4◦C on a rocking platform. To label thalamic calcification, sections were co-stained with AF647-RIS Imaging Reagent (BioVinc LLC, PV500101) diluted 1:500 in blocking buffer. Following overnight incubation, sections were washed 3 x 5 min with PBS and incubated for at least 2 h in fluorophore-labelled secondary antibodies (Donkey anti-rabbit AF647 (1:500, Invitrogen, A31573); Goat anti-chicken AF594 (1:500, Invitrogen, A32759)) diluted in blocking buffer. Sections were then washed 3 x 5 min with PBS before nuclei staining with DAPI (1:5000, ThermoFisher, #62248) for 5 min, washed in PBS and mounted onto glass slides with Fluorescence Mounting Medium (Dako), and stored at 4◦C in the dark until imaging.

For cryosections, spleens were dissected and immersion fixed in 4% PFA for 4h, before cryoprotection in 15% sucrose in PBS overnight and then 30% sucrose in PBS for 48 h. Spleens were then embedded in OCT, frozen and sectioned at 10μm using a ThermoFisher HM525NX cryostat. For staining, sections were rehydrated in Tris-buffered saline (TBS; 10mM Tris, 150mM NaCl, pH8.0) for 10 min and washed 3 x min in TBST (0.01% Tween-20 in TBS). Blocking buffer (3% BSA, 5% serum in TBST) was applied for 30 min at RT, before sections were incubated with primary antibodies: rat anti-CD169 (1:100, Biolegend, 142402) and rabbit anti-CD209B (1:150, Abcam, ab308457), diluted in TBST for 1.5 h at RT. Sections were washed 3 x 5 min TBST before incubation in secondary antibodies: donkey anti-rabbit AF647 (1:500, Invitrogen, A31573) and goat anti-rat AF594 (1:400, Invitrogen, A11007) diluted in TBST for 45 min protected from light. Sections were washed 3 x 5 min in PBS, stained with DAPI and mounted as above.

For bone staining the distal epiphysis of the femur and the fibular notch of the tibia were removed to expose the bone marrow. Tissues were fixed in 4% PFA for 24 hours at 4°C and washed in PBS prior to decalcification in 14% EDTA (pH 7.2) for 2 weeks at 4°C, cryoprotection by sequential incubation in 15% and 30% sucrose solutions, embedded in OCT and cryosectioned at 5 μm. For staining, sections were rehydrated with 1x TBS, blocked with 5% bovine serum albumin (BSA) + 10% normal donkey serum (NDS) + 10% normal goat serum (NGS), permeabilised with 0.1% Triton X-100 and stained with rat anti-mouse CD169 (Biolegend, Cat: 142402), rabbit anti-mouse F4/80 (Abcam, Cat: ab30042), rabbit anti-mouse CD71 (Invitrogen, Cat: MA5-48194) primary antibodies. After washing, sections were stained with secondary donkey anti-rat-AF647 (Invitrogen, Cat: A48272) and goat anti-rabbit-AF594 (Invitrogen, Cat: A32740) antibodies. Images were obtained using an Olympus FV3000. For TRAP staining, frozen tissue sections were washed with RO water and Acetate/Tartrate Buffer (150 mM Acetate, 6mM sodium tartrate, pH 5.0) then stained with ELF-97 (ThermoFisher Cat: E6588; 1:25 in Acetate/Tartrate Buffer,) for 10 mins, washed with RO water and mounted as above.

### Histochemistry

Tissues were fixed in 10% neutral buffered formalin and embedded in paraffin blocks. Sections were cut on a microtome (Leica) at 10 µm, laid on a 40°C water bath and collected on Superfrost Plus slides (Bio-strategy). Slides were incubated at 60 °C for 30 min before being de-waxed in xylene 2 x 5 min then rehydrated for 2 x 2 min washes in 100%, 95% and 75% ethanol, and washed in RO water for 5 min. Slides were stained with haematoxylin (Gill No 2, Sigma) for 30 seconds, rinsed under running tap water for 20 minutes and counterstained with Eosin Y (Sigma) for 40 seconds, dehydrated by 2 x 2 min washes in 70%, 95% and 100% ethanol, cleared 2 x 5 min in xylene and mounted with DPX mounting medium (Sigma). For immunohistochemistry, antigen retrieval was performed on dewaxed and rehydrated sections were incubated for 30 mins in 10 mM Tris/EDTA (pH 9) buffer maintained at 90-100°C using a microwave. Slides were cooled on ice for 10 min, incubated in 0.3% hydrogen peroxide (Sigma) for 15 min to block endogenous peroxidases and washed in TBS. Slides were incubated in 1% BSA in TBS for 30 min followed by incubation with primary antibody (anti-F4/80, 1:1000, rabbit, Abcam) for 1.5 hr in a humidified chamber. Slides were washed with TBS and HRP secondary reagent (anti-rabbit, Agilent) applied for 30 min. Slides were washed again with TBS and then incubated for 5 min with DAB Peroxidase substrate (Agilent). Slides were washed with TBS and counterstained with haematoxylin for 40 sec, then dehydrated and mounted as above. Imaging was performed on an Olympus BX50 microscope, using brightfield settings.

### Bone MicroCT

MicroCT was performed on formalin-fixed samples using a Skyscan 1272 desktop MicroCT (Bruker, Belgium). A 7 um voxel size was achieved using 4×4 camera binning and the following parameters: 70 kV voltage, 142 uA current, 1871 ms exposure time, averaging of 2, Al 0.5 mm filter, 0.5 degree rotation step and 360 degree rotation. The data was reconstructed with an FDK algorithm using NRecon 2.2.0.6 (Bruker, Belgium), and visualised with CTVox 3.3.1 (Bruker, Belgium). Volumetric analysis of the trabecular region of interest included bone volume percentage (BV/TV), trabecular thickness (Tb.Th), bone surface density (BS/TV), trabecular separation (Tb.Sp), and trabecular number (Tb.N). It was performed using CTAn 1.20.8.0 (Bruker, Belgium).

## Acknowledgements

The generation of the mice was funded originally by a grant from the Medical Research Council (MRC) UK grant MR/M019969/1 to DAH. This work was supported by core support and direct funding from The Mater Foundation and NHMRC Investigator Grant #2007850 to DAH. We acknowledge input and expertise from the Biological Resources facility and the Preclinical Imaging, Microscopy, Histology and Flow Cytometry facilities of the Translational Research Institute (TRI). TRI is supported by the Australian Government.

## References

1. Chitu, V., and Stanley, E.R. (2017). Regulation of Embryonic and Postnatal Development by the CSF-1 Receptor. Curr Top Dev Biol 123, 229–275. 10.1016/bs.ctdb.2016.10.004.

2. Hume, D.A., Caruso, M., Ferrari-Cestari, M., Summers, K.M., Pridans, C., and Irvine, K.M. (2020). Phenotypic impacts of CSF1R deficiencies in humans and model organisms. J Leukoc Biol 107, 205–219. 10.1002/JLB.MR0519-143R.

3. Stanley, E.R., and Chitu, V. (2014). CSF-1 receptor signaling in myeloid cells. Cold Spring Harb Perspect Biol 6. 10.1101/cshperspect.a021857.

4. Keshvari, S., Caruso, M., Teakle, N., Batoon, L., Sehgal, A., Patkar, O.L., Ferrari-Cestari, M., Snell, C.E., Chen, C., Stevenson, A., et al. (2021). CSF1R-dependent macrophages control postnatal somatic growth and organ maturation. PLoS Genet 17, e1009605. 10.1371/journal.pgen.1009605.

5. Pridans, C., Raper, A., David, G.M., Alves, J., Sauter, K.A., Lefevre, L., Regan, T., Grabert, K., Meek, S., Sutherland, L., et al. (2018). Pleiotropic Impacts of Macrophage and Microglial Deficiency on Development in Rats with Targeted Mutation of the Csf1r Locus. J.Immunol. 201(9), 2683–2699.

6. Sehgal, A., Carter-Cusack, D., Keshvari, S., Patkar, O., Huang, S., Summers, K.M., Hume, D.A., and Irvine, K.M. (2023). Intraperitoneal transfer of wild-type bone marrow repopulates tissue macrophages in the Csf1r knockout rat without contributing to monocytopoiesis. Eur J Immunol 53, e2250312. 10.1002/eji.202250312.

7. Hume, D.A., Teakle, N., Keshvari, S., and Irvine, K.M. (2023). Macrophage deficiency in CSF1R-knockout rat embryos does not compromise placental or embryo development. J Leukoc Biol 114, 421–433. 10.1093/jleuko/qiad052.

8. Guo, L., Bertola, D.R., Takanohashi, A., Saito, A., Segawa, Y., Yokota, T., Ishibashi, S., Nishida, Y., Yamamoto, G.L., Franco, J., et al. (2019). Bi-allelic CSF1R Mutations Cause Skeletal Dysplasia of Dysosteosclerosis-Pyle Disease Spectrum and Degenerative Encephalopathy with Brain Malformation. Am J Hum Genet 104, 925–935. 10.1016/j.ajhg.2019.03.004.

9. Oosterhof, N., Chang, I.J., Karimiani, E.G., Kuil, L.E., Jensen, D.M., Daza, R., Young, E., Astle, L., van der Linde, H.C., Shivaram, G.M., et al. (2019). Homozygous Mutations in CSF1R Cause a Pediatric-Onset Leukoencephalopathy and Can Result in Congenital Absence of Microglia. Am J Hum Genet 104, 936–947. 10.1016/j.ajhg.2019.03.010.

10. Dai, X.M., Ryan, G.R., Hapel, A.J., Dominguez, M.G., Russell, R.G., Kapp, S., Sylvestre, V., and Stanley, E.R. (2002). Targeted disruption of the mouse colony-stimulating factor 1 receptor gene results in osteopetrosis, mononuclear phagocyte deficiency, increased primitive progenitor cell frequencies, and reproductive defects. Blood 99, 111–120.

11. Li, J., Chen, K., Zhu, L., and Pollard, J.W. (2006). Conditional deletion of the colony stimulating factor-1 receptor (c-fms proto-oncogene) in mice. Genesis 44, 328–335. 10.1002/dvg.20219.

12. Percin, G.I., Eitler, J., Kranz, A., Fu, J., Pollard, J.W., Naumann, R., and Waskow, C. (2018). CSF1R regulates the dendritic cell pool size in adult mice via embryo-derived tissue-resident macrophages. Nat Commun 9, 5279. 10.1038/s41467-018-07685-x.

13. Elmore, M.R., Najafi, A.R., Koike, M.A., Dagher, N.N., Spangenberg, E.E., Rice, R.A., Kitazawa, M., Matusow, B., Nguyen, H., West, B.L., and Green, K.N. (2014). Colony-stimulating factor 1 receptor signaling is necessary for microglia viability, unmasking a microglia progenitor cell in the adult brain. Neuron 82, 380–397. 10.1016/j.neuron.2014.02.040.

14. Bogunovic, M., Ginhoux, F., Helft, J., Shang, L., Hashimoto, D., Greter, M., Liu, K., Jakubzick, C., Ingersoll, M.A., Leboeuf, M., et al. (2009). Origin of the lamina propria dendritic cell network. Immunity 31, 513–525. 10.1016/j.immuni.2009.08.010.

15. Chitu, V., Gokhan, S., and Stanley, E.R. (2021). Modeling CSF-1 receptor deficiency diseases - how close are we? FEBS J 289, 5049–5073. 10.1111/febs.16085.

16. Dulski, J., Souza, J., Santos, M.L., and Wszolek, Z.K. (2023). Brain abnormalities, neurodegeneration, and dysosteosclerosis (BANDDOS): new cases, systematic literature review, and associations with CSF1R-ALSP. Orphanet J Rare Dis 18, 160. 10.1186/s13023-023-02772-9.

17. Guo, L., and Ikegawa, S. (2021). From HDLS to BANDDOS: fast-expanding phenotypic spectrum of disorders caused by mutations in CSF1R. J Hum Genet 66, 1139–1144. 10.1038/s10038-021-00942-w.

18. Papapetropoulos, S., Gelfand, J.M., Konno, T., Ikeuchi, T., Pontius, A., Meier, A., Foroutan, F., and Wszolek, Z.K. (2024). Clinical presentation and diagnosis of adult-onset leukoencephalopathy with axonal spheroids and pigmented glia: a literature analysis of case studies. Front Neurol 15, 1320663. 10.3389/fneur.2024.1320663.

19. Wu, J., Cheng, X., Ji, D., Niu, H., Yao, S., Lv, X., Wang, J., Li, Z., Zheng, H., Cao, Y., et al. (2024). The Phenotypic and Genotypic Spectrum of CSF1R-Related Disorder in China. Mov Disord 39, 798–813. 10.1002/mds.29764.

20. Pridans, C., Sauter, K.A., Baer, K., Kissel, H., and Hume, D.A. (2013). CSF1R mutations in hereditary diffuse leukoencephalopathy with spheroids are loss of function. Sci Rep 3, 3013. 10.1038/srep03013.

21. Rademakers, R., Baker, M., Nicholson, A.M., Rutherford, N.J., Finch, N., Soto-Ortolaza, A., Lash, J., Wider, C., Wojtas, A., DeJesus-Hernandez, M., et al. (2011). Mutations in the colony stimulating factor 1 receptor (CSF1R) gene cause hereditary diffuse leukoencephalopathy with spheroids. Nat Genet 44, 200–205. 10.1038/ng.1027.

22. Munro, D.A.D., Bradford, B.M., Mariani, S.A., Hampton, D.W., Vink, C.S.S., Chandran, S., Hume, D.A., Pridans, C., and Priller, J. (2020). CNS macrophages differentially rely on an intronic Csf1r enhancer for their development. Development 147, dev194449.

23. Stables, J., Green, E.K., Sehgal, A., Patkar, O.L., Keshvari, S., Taylor, I., Ashcroft, M.E., Grabert, K., Wollscheid-Lengeling, E., Szymkowiak, S., et al. (2022). A kinase-dead Csf1r mutation associated with adult-onset leukoencephalopathy has a dominant inhibitory impact on CSF1R signalling. Development 149, dev200237. 10.1242/dev.200237.

24. Sasmono, R.T., Oceandy, D., Pollard, J.W., Tong, W., Pavli, P., Wainwright, B.J., Ostrowski, M.C., Himes, S.R., and Hume, D.A. (2003). A macrophage colony-stimulating factor receptor-green fluorescent protein transgene is expressed throughout the mononuclear phagocyte system of the mouse. Blood 101, 1155–1163. 10.1182/blood-2002-02-0569.

25. Helft, J., Bottcher, J., Chakravarty, P., Zelenay, S., Huotari, J., Schraml, B.U., Goubau, D., and Reis eSousa, C. (2015). GM-CSF Mouse Bone Marrow Cultures Comprise a Heterogeneous Population of CD11c(+)MHCII(+) Macrophages and Dendritic Cells. Immunity 42, 1197–1211. 10.1016/j.immuni.2015.05.018.

26. Carter-Cusack, D., Huang, S., Keshvari, S., Patkar, O., Sehgal, A., Allavena, R., Byrne, R.A.J., Morgan, B.P., Bush, S.J., Summers, K.M., et al. (2025). Wild-type bone marrow cells repopulate tissue resident macrophages and reverse the impacts of homozygous CSF1R mutation. PLoS Genet 21, e1011525. 10.1371/journal.pgen.1011525.

27. Bartocci, A., Mastrogiannis, D.S., Migliorati, G., Stockert, R.J., Wolkoff, A.W., and Stanley, E.R. (1987). Macrophages specifically regulate the concentration of their own growth factor in the circulation. Proc Natl Acad Sci U S A 84, 6179–6183. 10.1073/pnas.84.17.6179.

28. Sierro, F., Evrard, M., Rizzetto, S., Melino, M., Mitchell, A.J., Florido, M., Beattie, L., Walters, S.B., Tay, S.S., Lu, B., et al. (2017). A Liver Capsular Network of Monocyte-Derived Macrophages Restricts Hepatic Dissemination of Intraperitoneal Bacteria by Neutrophil Recruitment. Immunity 47, 374–388 e376. 10.1016/j.immuni.2017.07.018.

29. Hume, D.A., Robinson, A.P., MacPherson, G.G., and Gordon, S. (1983). The mononuclear phagocyte system of the mouse defined by immunohistochemical localization of antigen F4/80. Relationship between macrophages, Langerhans cells, reticular cells, and dendritic cells in lymphoid and hematopoietic organs. J Exp Med 158, 1522–1536. 10.1084/jem.158.5.1522.

30. Hohsfield, L.A., Kim, S.J., Barahona, R.A., Henningfield, C.M., Mansour, K., Vallejo, K.D., Tsourmas, K.I., Kwang, N.E., Ghorbanian, Y., Angulo, J.A.A., et al. (2025). Identification of the velum interpositum as a meningeal-CNS route for myeloid cell trafficking into the brain. Neuron. 10.1016/j.neuron.2025.05.004.

31. Erblich, B., Zhu, L., Etgen, A.M., Dobrenis, K., and Pollard, J.W. (2011). Absence of colony stimulation factor-1 receptor results in loss of microglia, disrupted brain development and olfactory deficits. PLoS One 6, e26317. 10.1371/journal.pone.0026317.

32. O’Connell, K.E., Mikkola, A.M., Stepanek, A.M., Vernet, A., Hall, C.D., Sun, C.C., Yildirim, E., Staropoli, J.F., Lee, J.T., and Brown, D.E. (2015). Practical murine hematopathology: a comparative review and implications for research. Comp Med 65, 96–113.

33. Grabert, K., Sehgal, A., Irvine, K.M., Wollscheid-Lengeling, E., Ozdemir, D.D., Stables, J., Luke, G.A., Ryan, M.D., Adamson, A., Humphreys, N.E., et al. (2020). A Transgenic Line That Reports CSF1R Protein Expression Provides a Definitive Marker for the Mouse Mononuclear Phagocyte System. J Immunol 205, 3154–3166. 10.4049/jimmunol.2000835.

34. Trzebanski, S., Kim, J.S., Larossi, N., Raanan, A., Kancheva, D., Bastos, J., Haddad, M., Solomon, A., Sivan, E., Aizik, D., et al. (2024). Classical monocyte ontogeny dictates their functions and fates as tissue macrophages. Immunity 57, 1225–1242 e1226. 10.1016/j.immuni.2024.04.019.

35. Chang, M.K., Raggatt, L.J., Alexander, K.A., Kuliwaba, J.S., Fazzalari, N.L., Schroder, K., Maylin, E.R., Ripoll, V.M., Hume, D.A., and Pettit, A.R. (2008). Osteal tissue macrophages are intercalated throughout human and mouse bone lining tissues and regulate osteoblast function in vitro and in vivo. J Immunol 181, 1232–1244. 10.4049/jimmunol.181.2.1232.

36. Batoon, L., Millard, S.M., Wullschleger, M.E., Preda, C., Wu, A.C., Kaur, S., Tseng, H.W., Hume, D.A., Levesque, J.P., Raggatt, L.J., and Pettit, A.R. (2019). CD169(+) macrophages are critical for osteoblast maintenance and promote intramembranous and endochondral ossification during bone repair. Biomaterials 196, 51–66. 10.1016/j.biomaterials.2017.10.033.

37. Kaur, S., Raggatt, L.J., Batoon, L., Hume, D.A., Levesque, J.P., and Pettit, A.R. (2017). Role of bone marrow macrophages in controlling homeostasis and repair in bone and bone marrow niches. Semin Cell Dev Biol 61, 12–21. 10.1016/j.semcdb.2016.08.009.

38. Kaur, S., Raggatt, L.J., Millard, S.M., Wu, A.C., Batoon, L., Jacobsen, R.N., Winkler, I.G., MacDonald, K.P., Perkins, A.C., Hume, D.A., et al. (2018). Self-repopulating recipient bone marrow resident macrophages promote long-term hematopoietic stem cell engraftment. Blood 132, 735–749. 10.1182/blood-2018-01-829663.

39. Millard, S.M., Heng, O., Opperman, K.S., Sehgal, A., Irvine, K.M., Kaur, S., Sandrock, C.J., Wu, A.C., Magor, G.W., Batoon, L., et al. (2021). Fragmentation of tissue-resident macrophages during isolation confounds analysis of single-cell preparations from mouse hematopoietic tissues. Cell Rep 37, 110058. 10.1016/j.celrep.2021.110058.

40. Pop, R., Shearstone, J.R., Shen, Q., Liu, Y., Hallstrom, K., Koulnis, M., Gribnau, J., and Socolovsky, M. (2010). A key commitment step in erythropoiesis is synchronized with the cell cycle clock through mutual inhibition between PU.1 and S-phase progression. PLoS Biol 8. 10.1371/journal.pbio.1000484.

41. Kang, Y.S., Yamazaki, S., Iyoda, T., Pack, M., Bruening, S.A., Kim, J.Y., Takahara, K., Inaba, K., Steinman, R.M., and Park, C.G. (2003). SIGN-R1, a novel C-type lectin expressed by marginal zone macrophages in spleen, mediates uptake of the polysaccharide dextran. Int Immunol 15, 177–186. 10.1093/intimm/dxg019.

42. Kang, Y.A., Paik, H., Zhang, S.Y., Chen, J.J., Olson, O.C., Mitchell, C.A., Collins, A., Swann, J.W., Warr, M.R., Fan, R., and Passegue, E. (2023). Secretory MPP3 reinforce myeloid differentiation trajectory and amplify myeloid cell production. J Exp Med 220. 10.1084/jem.20230088.

43. Pietras, E.M., Reynaud, D., Kang, Y.A., Carlin, D., Calero-Nieto, F.J., Leavitt, A.D., Stuart, J.M., Gottgens, B., and Passegue, E. (2015). Functionally Distinct Subsets of Lineage-Biased Multipotent Progenitors Control Blood Production in Normal and Regenerative Conditions. Cell Stem Cell 17, 35–46. 10.1016/j.stem.2015.05.003.

44. Chitu, V., Gokhan, S., Nandi, S., Mehler, M.F., and Stanley, E.R. (2016). Emerging Roles for CSF-1 Receptor and its Ligands in the Nervous System. Trends Neurosci 39, 378–393. 10.1016/j.tins.2016.03.005.

45. Stables, J., Paal, R., Bradford, B.M., Carter-Cusack, D., Taylor, I., Pridans, C., Khan, N., Woodruff, T.M., Irvine, K.M., Summers, K.M., et al. (2024). The effect of a dominant kinase-dead Csf1r mutation associated with adult-onset leukoencephalopathy on brain development and neuropathology. . Neurobiol. Disease *In press*.

46. Rojo, R., Raper, A., Ozdemir, D.D., Lefevre, L., Grabert, K., Wollscheid-Lengeling, E., Bradford, B., Caruso, M., Gazova, I., Sanchez, A., et al. (2019). Deletion of a Csf1r enhancer selectively impacts CSF1R expression and development of tissue macrophage populations. Nat Commun 10, 3215. 10.1038/s41467-019-11053-8.

47. Lawrence, A.R., Canzi, A., Bridlance, C., Olivie, N., Lansonneur, C., Catale, C., Pizzamiglio, L., Kloeckner, B., Silvin, A., Munro, D.A.D., et al. (2024). Microglia maintain structural integrity during fetal brain morphogenesis. Cell 187, 962–980 e919. 10.1016/j.cell.2024.01.012.

48. Hume, D.A. (2023). Fate-mapping studies in inbred mice: A model for understanding macrophage development and homeostasis? Eur J Immunol 53, e2250242. 10.1002/eji.202250242.

49. Cortes-Hernandez, J., Fossati-Jimack, L., Petry, F., Loos, M., Izui, S., Walport, M.J., Cook, H.T., and Botto, M. (2004). Restoration of C1q levels by bone marrow transplantation attenuates autoimmune disease associated with C1q deficiency in mice. Eur J Immunol 34, 3713–3722. 10.1002/eji.200425616.

50. Gallerand, A., Han, J., Ivanov, S., and Randolph, G.J. (2024). Mouse and human macrophages and their roles in cardiovascular health and disease. Nat Cardiovasc Res 3, 1424–1437. 10.1038/s44161-024-00580-3.

51. Mass, E., Nimmerjahn, F., Kierdorf, K., and Schlitzer, A. (2023). Tissue-specific macrophages: how they develop and choreograph tissue biology. Nat Rev Immunol 23, 563–579. 10.1038/s41577-023-00848-y.

52. Lazarov, T., Juarez-Carreno, S., Cox, N., and Geissmann, F. (2023). Physiology and diseases of tissue-resident macrophages. Nature 618, 698–707. 10.1038/s41586-023-06002-x.

53. Witmer-Pack, M.D., Hughes, D.A., Schuler, G., Lawson, L., McWilliam, A., Inaba, K., Steinman, R.M., and Gordon, S. (1993). Identification of macrophages and dendritic cells in the osteopetrotic (op/op) mouse. J Cell Sci 104 (Pt 4), 1021–1029. 10.1242/jcs.104.4.1021.

54. Pirgova, G., Chauveau, A., MacLean, A.J., Cyster, J.G., and Arnon, T.I. (2020). Marginal zone SIGN-R1(+) macrophages are essential for the maturation of germinal center B cells in the spleen. Proc Natl Acad Sci U S A 117, 12295–12305. 10.1073/pnas.1921673117.

55. Steiniger, B.S. (2015). Human spleen microanatomy: why mice do not suffice. Immunology 145, 334–346. 10.1111/imm.12469.

56. Ginhoux, F., Greter, M., Leboeuf, M., Nandi, S., See, P., Gokhan, S., Mehler, M.F., Conway, S.J., Ng, L.G., Stanley, E.R., et al. (2010). Fate mapping analysis reveals that adult microglia derive from primitive macrophages. Science 330, 841–845. 10.1126/science.1194637.

57. Nandi, S., Gokhan, S., Dai, X.M., Wei, S., Enikolopov, G., Lin, H., Mehler, M.F., and Stanley, E.R. (2012). The CSF-1 receptor ligands IL-34 and CSF-1 exhibit distinct developmental brain expression patterns and regulate neural progenitor cell maintenance and maturation. Dev Biol 367, 100–113. 10.1016/j.ydbio.2012.03.026.

58. Patkar, O.L., Caruso, M., Teakle, N., Keshvari, S., Bush, S.J., Pridans, C., Belmer, A., Summers, K.M., Irvine, K.M., and Hume, D.A. (2021). Analysis of homozygous and heterozygous Csf1r knockout in the rat as a model for understanding microglial function in brain development and the impacts of human CSF1R mutations. Neurobiol Dis 151, 105268. 10.1016/j.nbd.2021.105268.

59. Chadarevian, J.P., Hasselmann, J., Escobar, A., Lahian, A., Lim, T.E., Capocchi, J., Tu, C., Nguyen, J., Kiani Shabestari, S., Carmen-Jones, W., et al. (2024). Therapeutic potential of human microglial transplantation in a chimeric model of CSF1R-related leukoencephalopathy. . Neuron 112, 2686–2707.

60. Munro, D.A.D., Bestard-Cuche, N., McQuade, A., Chagnot, A., Kiani Shabestari, S., Chadarevian, J.P., Maheshwari, U., Szymkowiak, S., Morris, K., Mohammad, M., et al. (2024). Microglia protect against age-associated brain pathologies. . Neuron 112, 2732–2748.

61. Akcora, D., Huynh, D., Lightowler, S., Germann, M., Robine, S., de May, J.R., Pollard, J.W., Stanley, E.R., Malaterre, J., and Ramsay, R.G. (2013). The CSF-1 receptor fashions the intestinal stem cell niche. Stem Cell Res 10, 203–212. 10.1016/j.scr.2012.12.001.

62. Huynh, D., Akcora, D., Malaterre, J., Chan, C.K., Dai, X.M., Bertoncello, I., Stanley, E.R., and Ramsay, R.G. (2013). CSF-1 receptor-dependent colon development, homeostasis and inflammatory stress response. PLoS One 8, e56951. 10.1371/journal.pone.0056951.

63. Huynh, D., Dai, X.M., Nandi, S., Lightowler, S., Trivett, M., Chan, C.K., Bertoncello, I., Ramsay, R.G., and Stanley, E.R. (2009). Colony stimulating factor-1 dependence of paneth cell development in the mouse small intestine. Gastroenterology 137, 136-144, 144 e131–133. 10.1053/j.gastro.2009.03.004.

64. Sehgal, A., Donaldson, D.S., Pridans, C., Sauter, K.A., Hume, D.A., and Mabbott, N.A. (2018). The role of CSF1R-dependent macrophages in control of the intestinal stem-cell niche. Nat Commun 9, 1272. 10.1038/s41467-018-03638-6.

65. Bain, C.C., Bravo-Blas, A., Scott, C.L., Perdiguero, E.G., Geissmann, F., Henri, S., Malissen, B., Osborne, L.C., Artis, D., and Mowat, A.M. (2014). Constant replenishment from circulating monocytes maintains the macrophage pool in the intestine of adult mice. Nat Immunol 15, 929–937. 10.1038/ni.2967.

66. Hernandez-Barrientos, D., Pelayo, R., and Mayani, H. (2023). The hematopoietic microenvironment: a network of niches for the development of all blood cell lineages. J Leukoc Biol 114, 404–420. 10.1093/jleuko/qiad075.

67. Pinho, S., Wei, Q., Maryanovich, M., Zhang, D., Balandran, J.C., Pierce, H., Nakahara, F., Di Staulo, A., Bartholdy, B.A., Xu, J., et al. (2022). VCAM1 confers innate immune tolerance on haematopoietic and leukaemic stem cells. Nat Cell Biol 24, 290–298. 10.1038/s41556-022-00849-4.

68. Wei, Q., Boulais, P.E., Zhang, D., Pinho, S., Tanaka, M., and Frenette, P.S. (2019). Maea expressed by macrophages, but not erythroblasts, maintains postnatal murine bone marrow erythroblastic islands. Blood 133, 1222–1232. 10.1182/blood-2018-11-888180.

69. Kayvanjoo, A.H., Splichalova, I., Bejarano, D.A., Huang, H., Mauel, K., Makdissi, N., Heider, D., Tew, H.M., Balzer, N.R., Greto, E., et al. (2024). Fetal liver macrophages contribute to the hematopoietic stem cell niche by controlling granulopoiesis. Elife 13. 10.7554/eLife.86493.

70. Romano, L., Seu, K.G., Papoin, J., Muench, D.E., Konstantinidis, D., Olsson, A., Schlum, K., Chetal, K., Chasis, J.A., Mohandas, N., et al. (2022). Erythroblastic islands foster granulopoiesis in parallel to terminal erythropoiesis. Blood 140, 1621–1634. 10.1182/blood.2022015724.

71. Sarrazin, S., Mossadegh-Keller, N., Fukao, T., Aziz, A., Mourcin, F., Vanhille, L., Kelly Modis, L., Kastner, P., Chan, S., Duprez, E., et al. (2009). MafB restricts M-CSF-dependent myeloid commitment divisions of hematopoietic stem cells. Cell 138, 300–313. 10.1016/j.cell.2009.04.057.

72. Mossadegh-Keller, N., Sarrazin, S., Kandalla, P.K., Espinosa, L., Stanley, E.R., Nutt, S.L., Moore, J., and Sieweke, M.H. (2013). M-CSF instructs myeloid lineage fate in single haematopoietic stem cells. Nature 497, 239–243. 10.1038/nature12026.

73. Hume, D.A., Millard, S., and Pettit, A.R. (2023). Macrophage biology in the single cell era: facts and artefacts. Blood 142, 1339–1347.

74. Chow, A., Huggins, M., Ahmed, J., Hashimoto, D., Lucas, D., Kunisaki, Y., Pinho, S., Leboeuf, M., Noizat, C., van Rooijen, N., et al. (2013). CD169(+) macrophages provide a niche promoting erythropoiesis under homeostasis and stress. Nat Med 19, 429–436. 10.1038/nm.3057.

75. Gow, D.J., Sauter, K.A., Pridans, C., Moffat, L., Sehgal, A., Stutchfield, B.M., Raza, S., Beard, P.M., Tsai, Y.T., Bainbridge, G., et al. (2014). Characterisation of a novel Fc conjugate of macrophage colony-stimulating factor. Mol Ther 22, 1580–1592. 10.1038/mt.2014.112.

76. Keshvari, S., Masson, J.J.R., Ferrari-Cestari, M., Bodea, L.G., Nooru-Mohamed, F., Tse, B.W.C., Sokolowski, K.A., Batoon, L., Patkar, O.L., Sullivan, M.A., et al. (2024). Reversible expansion of tissue macrophages in response to macrophage colony-stimulating factor (CSF1) transforms systemic lipid and carbohydrate metabolism. Am J Physiol Endocrinol Metab 326, E149–E165. 10.1152/ajpendo.00347.2023.

77. Louis, C., Cook, A.D., Lacey, D., Fleetwood, A.J., Vlahos, R., Anderson, G.P., and Hamilton, J.A. (2015). Specific Contributions of CSF-1 and GM-CSF to the Dynamics of the Mononuclear Phagocyte System. J Immunol 195, 134–144. 10.4049/jimmunol.1500369.

78. MacDonald, K.P., Palmer, J.S., Cronau, S., Seppanen, E., Olver, S., Raffelt, N.C., Kuns, R., Pettit, A.R., Clouston, A., Wainwright, B., et al. (2010). An antibody against the colony-stimulating factor 1 receptor depletes the resident subset of monocytes and tissue- and tumor-associated macrophages but does not inhibit inflammation. Blood 116, 3955–3963. 10.1182/blood-2010-02-266296.

79. Hashimoto, D., Chow, A., Greter, M., Saenger, Y., Kwan, W.H., Leboeuf, M., Ginhoux, F., Ochando, J.C., Kunisaki, Y., van Rooijen, N., et al. (2011). Pretransplant CSF-1 therapy expands recipient macrophages and ameliorates GVHD after allogeneic hematopoietic cell transplantation. J Exp Med 208, 1069–1082. 10.1084/jem.20101709.

80. Emoto, T., Lu, J., Sivasubramaniyam, T., Maan, H., Khan, A.B., Abow, A.A., Schroer, S.A., Hyduk, S.J., Althagafi, M.G., McKee, T.D., et al. (2022). Colony stimulating factor-1 producing endothelial cells and mesenchymal stromal cells maintain monocytes within a perivascular bone marrow niche. Immunity 55, 862–878 e868. 10.1016/j.immuni.2022.04.005.

81. Thierry, G.R., Baudon, E.M., Bijnen, M., Bellomo, A., Lagueyrie, M., Mondor, I., Simonnet, L., Carrette, F., Fenouil, R., Keshvari, S., et al. (2024). Non-classical monocytes scavenge the growth factor CSF1 from endothelial cells in the peripheral vascular tree to ensure survival and homeostasis. Immunity 57, 2108–2121 e2106. 10.1016/j.immuni.2024.07.005.

82. Zhang, J., Wu, Q., Johnson, C.B., Pham, G., Kinder, J.M., Olsson, A., Slaughter, A., May, M., Weinhaus, B., D’Alessandro, A., et al. (2021). In situ mapping identifies distinct vascular niches for myelopoiesis. Nature 590, 457–462. 10.1038/s41586-021-03201-2.

83. Greter, M., Helft, J., Chow, A., Hashimoto, D., Mortha, A., Agudo-Cantero, J., Bogunovic, M., Gautier, E.L., Miller, J., Leboeuf, M., et al. (2012). GM-CSF controls nonlymphoid tissue dendritic cell homeostasis but is dispensable for the differentiation of inflammatory dendritic cells. Immunity 36, 1031–1046. 10.1016/j.immuni.2012.03.027.

84. Sudo, T., Nishikawa, S., Ogawa, M., Kataoka, H., Ohno, N., Izawa, A., Hayashi, S., and Nishikawa, S. (1995). Functional hierarchy of c-kit and c-fms in intramarrow production of CFU-M. Oncogene 11, 2469–2476.

85. Momenilandi, M., Levy, R., Sobrino, S., Li, J., Lagresle-Peyrou, C., Esmaeilzadeh, H., Fayand, A., Le Floc’h, C., Guerin, A., Della Mina, E., et al. (2024). FLT3L governs the development of partially overlapping hematopoietic lineages in humans and mice. Cell 187, 2817–2837 e2831. 10.1016/j.cell.2024.04.009.

86. Brasel, K., McKenna, H.J., Charrier, K., Morrissey, P.J., Williams, D.E., and Lyman, S.D. (1997). Flt3 ligand synergizes with granulocyte-macrophage colony-stimulating factor or granulocyte colony-stimulating factor to mobilize hematopoietic progenitor cells into the peripheral blood of mice. Blood 90, 3781–3788.

87. Brasel, K., McKenna, H.J., Morrissey, P.J., Charrier, K., Morris, A.E., Lee, C.C., Williams, D.E., and Lyman, S.D. (1996). Hematologic effects of flt3 ligand in vivo in mice. Blood 88, 2004–2012.

88. Lean, J.M., Fuller, K., and Chambers, T.J. (2001). FLT3 ligand can substitute for macrophage colony-stimulating factor in support of osteoclast differentiation and function. Blood 98, 2707–2713. 10.1182/blood.v98.9.2707.

89. Jacquelin, S., Licata, F., Dorgham, K., Hermand, P., Poupel, L., Guyon, E., Deterre, P., Hume, D.A., Combadiere, C., and Boissonnas, A. (2013). CX3CR1 reduces Ly6Chigh-monocyte motility within and release from the bone marrow after chemotherapy in mice. Blood 122, 674–683. 10.1182/blood-2013-01-480749.

90. Guilliams, M., Mildner, A., and Yona, S. (2018). Developmental and Functional Heterogeneity of Monocytes. Immunity 49, 595–613. 10.1016/j.immuni.2018.10.005.

91. Yona, S., Kim, K.W., Wolf, Y., Mildner, A., Varol, D., Breker, M., Strauss-Ayali, D., Viukov, S., Guilliams, M., Misharin, A., et al. (2013). Fate mapping reveals origins and dynamics of monocytes and tissue macrophages under homeostasis. Immunity 38, 79–91. 10.1016/j.immuni.2012.12.001.

92. De Filippo, K., and Rankin, S.M. (2018). CXCR4, the master regulator of neutrophil trafficking in homeostasis and disease. Eur J Clin Invest 48 Suppl 2, e12949. 10.1111/eci.12949.

93. Rankin, S.M. (2010). The bone marrow: a site of neutrophil clearance. J Leukoc Biol 88, 241–251. 10.1189/jlb.0210112.

94. Klei, T.R., Meinderts, S.M., van den Berg, T.K., and van Bruggen, R. (2017). From the Cradle to the Grave: The Role of Macrophages in Erythropoiesis and Erythrophagocytosis. Front Immunol 8, 73. 10.3389/fimmu.2017.00073.

95. Zriwil, A., Boiers, C., Wittmann, L., Green, J.C., Woll, P.S., Jacobsen, S.E., and Sitnicka, E. (2016). Macrophage colony-stimulating factor receptor marks and regulates a fetal myeloid-primed B-cell progenitor in mice. Blood 128, 217–226. 10.1182/blood-2016-01-693887.

96. Corder, K.M., Hoffman, J.M., Sogorovic, A., and Austad, S.N. (2023). Behavioral comparison of the C57BL/6 inbred mouse strain and their CB6F1 siblings. Behav Processes 207, 104836. 10.1016/j.beproc.2023.104836.

97. Soni, N., Hohsfield, L.A., Tran, K.M., Kawauchi, S., Walker, A., Javonillo, D., Phan, J., Matheos, D., Da Cunha, C., Uyar, A., et al. (2024). Genetic diversity promotes resilience in a mouse model of Alzheimer’s disease. Alzheimers Dement 20, 2794–2816. 10.1002/alz.13753.

98. Tuttle, A.H., Philip, V.M., Chesler, E.J., and Mogil, J.S. (2018). Comparing phenotypic variation between inbred and outbred mice. Nat Methods 15, 994–996. 10.1038/s41592-018-0224-7.

99. Hoeksema, M.A., Shen, Z., Holtman, I.R., Zheng, A., Spann, N.J., Cobo, I., Gymrek, M., and Glass, C.K. (2021). Mechanisms underlying divergent responses of genetically distinct macrophages to IL-4. Sci Adv 7. 10.1126/sciadv.abf9808.

100. Link, V.M., Duttke, S.H., Chun, H.B., Holtman, I.R., Westin, E., Hoeksema, M.A., Abe, Y., Skola, D., Romanoski, C.E., Tao, J., et al. (2018). Analysis of Genetically Diverse Macrophages Reveals Local and Domain-wide Mechanisms that Control Transcription Factor Binding and Function. Cell 173, 1796–1809 e1717. 10.1016/j.cell.2018.04.018.

101. Taylor, I. et al. (2025). Repopulation of the brain with microglia-like cells following intraperitoneal bone marrow cell transfer in microglia-deficient mice. bioRxiv 2025, 2025.2001.2016.633478.

102. Hofer, M.J., Finger, C., and Pagenstecher, A. (2005). Molecular genotyping of the murine H-2K MHC class I allele. J Immunol Methods 302, 168–171. 10.1016/j.jim.2005.04.001.

